# Pulcherrimin formation controls growth arrest of the *Bacillus subtilis* biofilm

**DOI:** 10.1101/570630

**Authors:** Sofia Arnaouteli, Daniel Matoz-Fernandez, Michael Porter, Margarita Kalamara, James Abbott, Cait E. MacPhee, Fordyce A. Davidson, Nicola R. Stanley-Wall

## Abstract

Biofilm formation by *Bacillus subtilis* is a communal process that culminates in the formation of architecturally complex multicellular communities. Here we reveal that the transition of the biofilm into a non-expanding phase constitutes a distinct step in the process of biofilm development. Using genetic analysis we show that *B. subtilis* strains lacking the ability to synthesize pulcherriminic acid form biofilms that sustain the expansion phase, thereby linking pulcherriminic acid to growth arrest. However, production of pulcherriminic acid is not sufficient to block expansion of the biofilm. It needs to be secreted into the extracellular environment where it chelates Fe^3+^ from the growth medium in a non-enzymatic reaction. Utilizing mathematical modelling and a series of experimental methodologies we show that when the level of freely available iron in the environment drops below a critical threshold, expansion of the biofilm stops. Bioinformatics analysis allows us to identify the genes required for pulcherriminic acid synthesis in other Firmicutes but the patchwork presence both within and across closely related species suggests loss of these genes through multiple independent recombination events. The seemingly counterintuitive self-restriction of growth led us to explore if there were any benefits associated pulcherriminic acid production. We identified that pulcherriminic acid producers can prevent invasion from neighbouring communities through the generation of an “iron free” zone thereby addressing the paradox of pulcherriminic acid production by *B. subtilis*.

**Significance:** Understanding the processes that underpin the mechanism of biofilm formation, dispersal, and inhibition are critical to allow exploitation and to understand how microbes thrive in the environment. Here, we reveal that the formation of an extracellular iron chelate restricts the expansion of a biofilm. The countering benefit to self-restriction of growth is protection of an environmental niche. These findings highlight the complex options and outcomes that bacteria need to balance in order to modulate their local environment to maximise colonisation, and therefore survival.

## Introduction

Biofilm formation is a survival strategy used by microorganisms to overcome adverse conditions. It is stimulated by diverse environmental signals that, for example, direct the production of the biofilm matrix (1). The extracellular polysaccharides, adhesins and matrix proteins that are synthesised by the bacterial population surround and attach to the cells (2). The biofilm matrix provides structure to the three-dimensional community and also fulfils many other functions (3) including the sequestration of water and nutrients (4), driving expansion across surfaces, conferring resistance to phage (5), and neutralising aminoglycoside antibiotics (6).

*Bacillus subtilis* is a Gram-positive, spore forming bacterium which is ubiquitous in the soil and rhizosphere environment. *B. subtilis* biofilms are of agricultural importance as they are linked with plant growth promotion and the provision of protection from pathogens when formed on plant root systems (7, 8). The mature *B. subtilis* biofilm manifests *in vitro* as an architecturally complex community (9) that is resistant to gas penetration (10) and exhibits extreme hydrophobicity (10, 11). The extracellular biofilm matrix needed for the assembly of the community comprises stable fibres formed by the secreted protein TasA (12–14), an exopolysaccharide (9), and a protein with dual functions in biofilm architecture and hydrophobicity called BslA (15–17).

The process of *B. subtilis* biofilm formation, in the laboratory, begins with the deposition of “founder” cells (18). These cells differentiate to undertake distinct roles (19–21). For example a sub-population of these cells transcribe the operons needed for biofilm matrix synthesis (22). Over time, the initial founding cell population divides and expands across the surface in an extracellular matrix dependent manner (18, 23, 24). However, after a period of 2-3 days, expansion of the biofilm stops. Here we find that days after the biofilm has stopped expanding, a proportion of metabolically active cells remain in the community. Thus, growth arrest is not due to sporulation of the entire population. Rather, we reveal that it is a distinct stage of biofilm formation by *B. subtilis*. Using a combination of genetics and mathematical modelling we connect synthesis of the extracellular iron chelator pulcherriminic acid, and the subsequent deposition of the iron chelate pulcherrimin, to the arrest of biofilm expansion. We determine that in the absence of pulcherriminic acid production, *B. subtilis* forms “ever expanding” biofilms. We uncover that pulcherriminic acid manipulates the microenvironment of the biofilm through depletion of iron in the resultant pulcherrimin deposit. Complete depletion of iron in the surrounding environment allows *B. subtilis* to defend its niche from neighbouring bacteria, whereas a partial depletion in high iron conditions allows *B. subtilis* to colonise a surface and gain access to nutrients. Taken together these findings highlight a new route by which a bacterial biofilm can optimise survival within a changing environment.

## Results

### Growth arrest is a distinct stage of biofilm formation

*B. subtilis* strain NCIB 3610 forms highly structured hydrophobic biofilms at the air-agar interface. We have noted that mature biofilms reach a finite size (Fig. 1A), where the footprint occupied by the community is largely independent of the area of nutrients provided. For example, the average footprints of 5-day old NCIB 3610 biofilms were ~5.4 cm^2^ and ~6.1 cm^2^ when formed on nutrient surfaces with areas ~176 cm^2^ and ~63.5 cm^2^, respectively (Fig. 1B, Fig. S1A). It is known that the genetic background of the bacterium (9) and the environmental conditions used for growth (18, 25) influence the area occupied by the biofilm. For example, using a softer agar (Fig. S1B) or depositing the equivalent number of initial cells in a larger volume (Fig S1C) increases the size of the biofilm footprint. However, regardless of experimental variations, the area colonised by the NCIB 3610 biofilm plateaus, despite the apparent availability of nutrients in the surrounding unoccupied surface.

**Figure 1:**
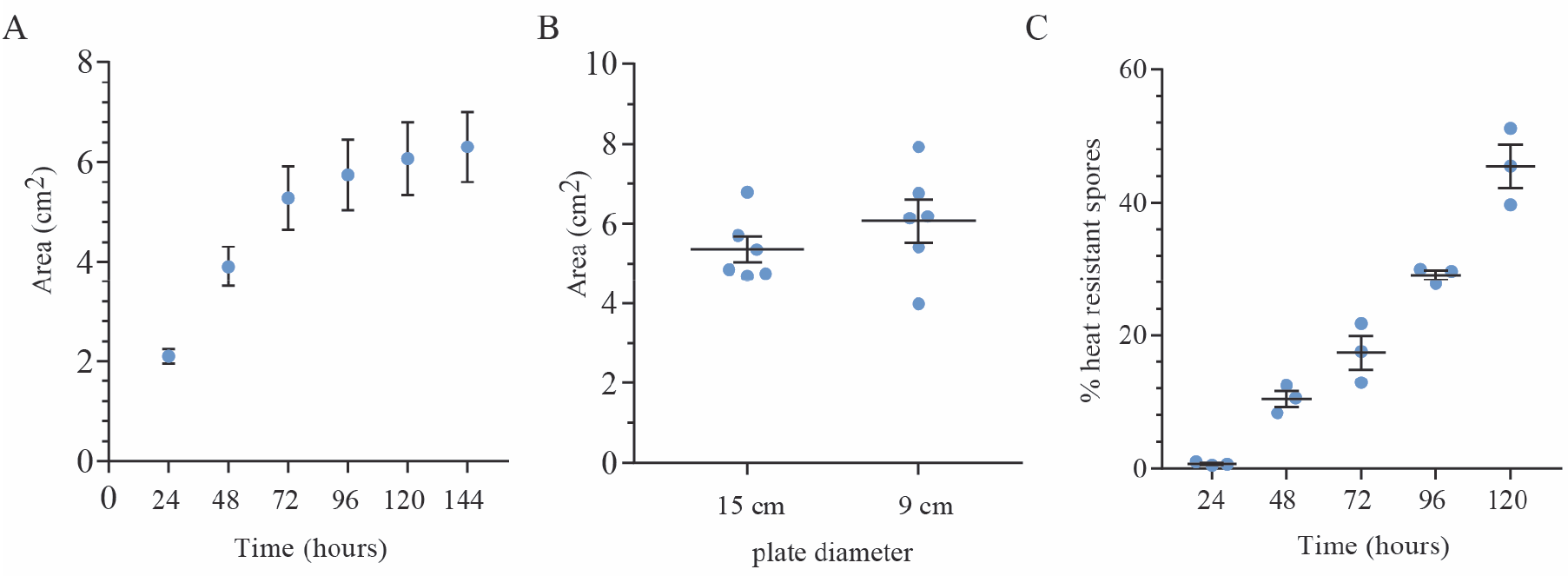
Growth arrest is a distinct stage of biofilm formation. **(A)** The footprint occupied by the NCIB 3610 biofilm was measured over time. The average of three biological repeats is presented, the error bars are the standard error of the mean; **(B)** The footprint of the NCIB 3610 biofilm was measured after 120 hours growth on agar plates with different diameters; **(C)** The number of heat resistant spores formed by NCIB 3610 was determined at the time points indicated. In all cases the biofilms were grown at 30°C on MSgg agar. For (B) and (C) the bar represents the average of the biological repeats that are presented as individual points, the error bars are the standard error of the mean.

*B. subtilis* is an endospore forming bacterium and the number of spores increases in the biofilm over time (21, 22). Therefore, a simple hypothesis to explain the arrest in biofilm expansion is that all of the cells in the biofilm have formed spores by 72 hours (the point at which the biofilm stops rapidly expanding (see Fig. 1A)), which would render the cells in the biofilm metabolically inactive. However, we found that while the number of heat resistant spores within the wild-type biofilm gradually increased over time, even at 120 hours the percentage of spores does not reach 100% of the total biofilm cell population (Fig. 1C). This leads to the conclusion that a population of cells in the wild-type 120-hour old biofilm remain metabolically active and therefore biofilm expansion must be restricted by some other process.

### Strains lacking pulcherriminic acid synthesis form “ever-expanding” biofilms

As the mature biofilm contains a significant proportion of cells that are not spores, the arrest in biofilm expansion cannot be due to metabolic inactivity. Therefore we explored other mechanisms. In biofilms formed by other species of bacteria, pigment production has been linked with biofilm morphology, cell activity, pathogenesis, and survival (26). The *B. subtilis* biofilm produces a secreted, brown pigment that is found in the agar underneath and surrounding the biomass (Fig S1A). We therefore explored if there was a link between production of the pigment and the arrest of biofilm expansion. Pulcherrimin is the only pigment known to be made by *B. subtilis* (27) and is produced when YvmC and CypX convert two tRNA-leucine molecules to pulcherriminic acid. Pulcherriminic acid is secreted from the cell where it binds Fe^3+^ in a non-enzymatic reaction (Fig. 2A) (28). The respective genes, *yvmC* and *cypX* (28), are co-located on the genome with the coding regions for a transcriptional regulator (PchR) (29) and a putative membrane bound transporter YvmA (30) (Fig. 2B).

**Figure 2:**
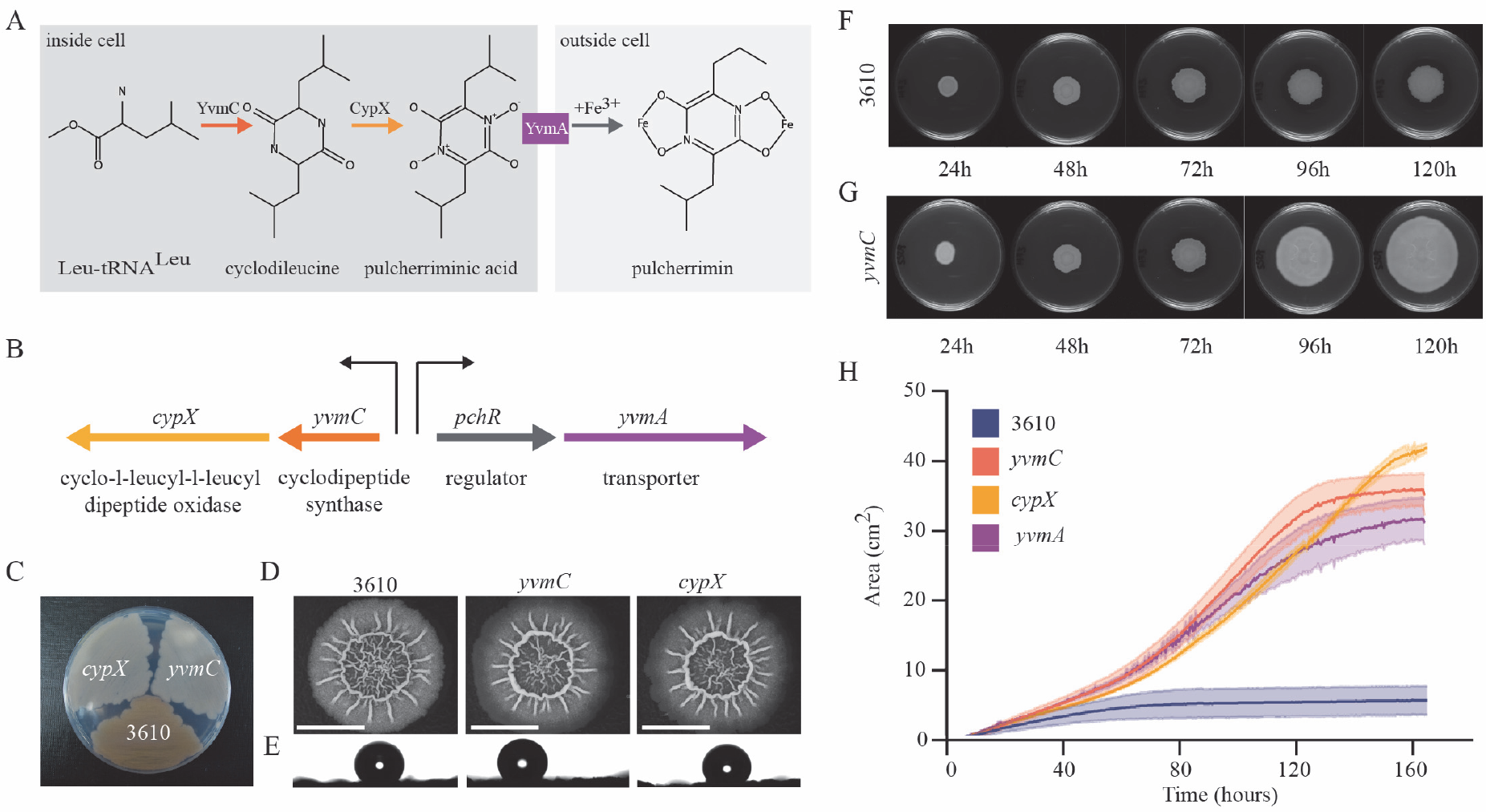
Pulcherrimin production restricts biofilm expansion. **(A)** Schematic depicting the steps involved in the formation of pulcherrimin; **(B)** Genomic organisation of the pulcherriminic acid biosynthesis cluster, the arrows represent promoters; **(C)** NCIB 3610 and the *cypX* (NRS5532) and *yvmC* (NRS5533) deletion strains after growth on MSgg agar for 24 hours at 37°C; **(D)** Image of biofilms formed by the strains detailed in (C); **(E)** a 5 μl water droplet placed on the biofilms depicted in (D) reveals hydrophobicity. For (D) and (E) the biofilms were grown on MSgg agar for 48 hours at 30°C; **(F, G)** Representative images from a time course of biofilm formation showing the area covered by the NCIB 3610 **(F)** and the *yvmC* mutant **(G)**; **(H)** biofilms formed by strains NCIB 3610, *yvmC* (NRS5533), *cypX* (NRS5532) and *yvmA* (NRS6248) were imaged every 20 minutes for 160 hours. The area occupied on the 9 cm diameter petri dish was calculated and plotted. The solid lines represent the average of 3 biological repeats and the shaded area the standard error of the mean. The samples in (F-H) were grown at 30°C on MSgg agar containing 50 μM FeCl_3_.

By introducing mutations in *yvmC* and *cypX* into the parental NCIB 3610 strain, we confirmed that the brown pigment was no longer produced (Fig. 2C). We therefore concluded that the pigment produced by the NCIB 3610 biofilm was pulcherrimin. We went on to examine if the *yvmC* and *cypX* deletions impacted biofilm formation. For the first 48 hrs of growth, the phenotypes of the wild-type and deletion mutant biofilms were virtually indistinguishable with respect to gross architecture (Fig 2D) and hydrophobicity (Fig. 2E). In contrast, at later time points, an obvious phenotypic difference manifested. As mentioned earlier, the area occupied by the NCIB 3610 biofilm stops increasing (Fig. 2F, 2H). In the absence of pulcherrimin formation, the biofilms formed by *yvmC* and *cypX* deletion strains continued to expand over time, almost occupying the entire substratum provided (Fig. 2G, 2H, S2A). The ability to keep colonising the surface was not due to a second site mutation in the genome as the *yvmC* and *cypX* deletion strains could be genetically complemented by provision of the *cypX* and *yvmC* genes under the control of their native promoter, at the ectopic *amyE* locus (Fig. S2A, S2B). These experiments reveal that production of pulcherriminic acid restricts the expansion of the *B. subtilis* biofilm community.

### Radial expansion of the pulcherrimin deficient strains

To explore how the pulcherriminic acid minus strains expand across the substratum, we assessed if there were differences in cell organisation at the outer (leading) edge of the biofilm using *in situ* confocal microscopy. To allow definition to be resolved in the densely packed bacterial community we mixed isogenic cells that constitutively expressed the green fluorescent protein with non-fluorescent cells (in a 1:5 ratio). We examined the outer biofilm edge at two time points: 24 hours and 120 hours. No detectable phenotypic differences between wild type and pulcherriminic acid deficient strains were observed at 24 hours of growth (Fig. 3A). The imaging revealed that the leading edge of the wild-type, *yvmC* and *cypX* deletion strains contained loops of cells that extended out from the body of the biofilm biomass. The loops protruding from the biofilm were located in what appeared to be a densely packed monolayer. The cells in the monolayer looked to be comprised of clonal lineages of cells as fluorescence was located in convoluted but discrete chains. In contrast, at 120 hours in the wild-type biofilm, the monolayer at the leading edge of the biofilm was absent and the cells were contained within a multilayer structure that consistently presented with a tight, compact steep edge in which chains or loops of cells could not be resolved. For the *cypX* and *yvmC* deletion strains, while certain zones of the 120 hour old biofilm edge also showed steep compact edges containing multiple layers of cells (Fig. S3), monolayers containing loops of cells that protruded from the body of the biofilm were maintained in many regions around the edge of the expanding community (Fig. 3A). These findings reveal a difference in the cell organisation of the pulcherriminic acid positive and negative strains and link the presence of a monolayer of cells at the periphery of the biofilm with expansion across the surface.

**Figure 3:**
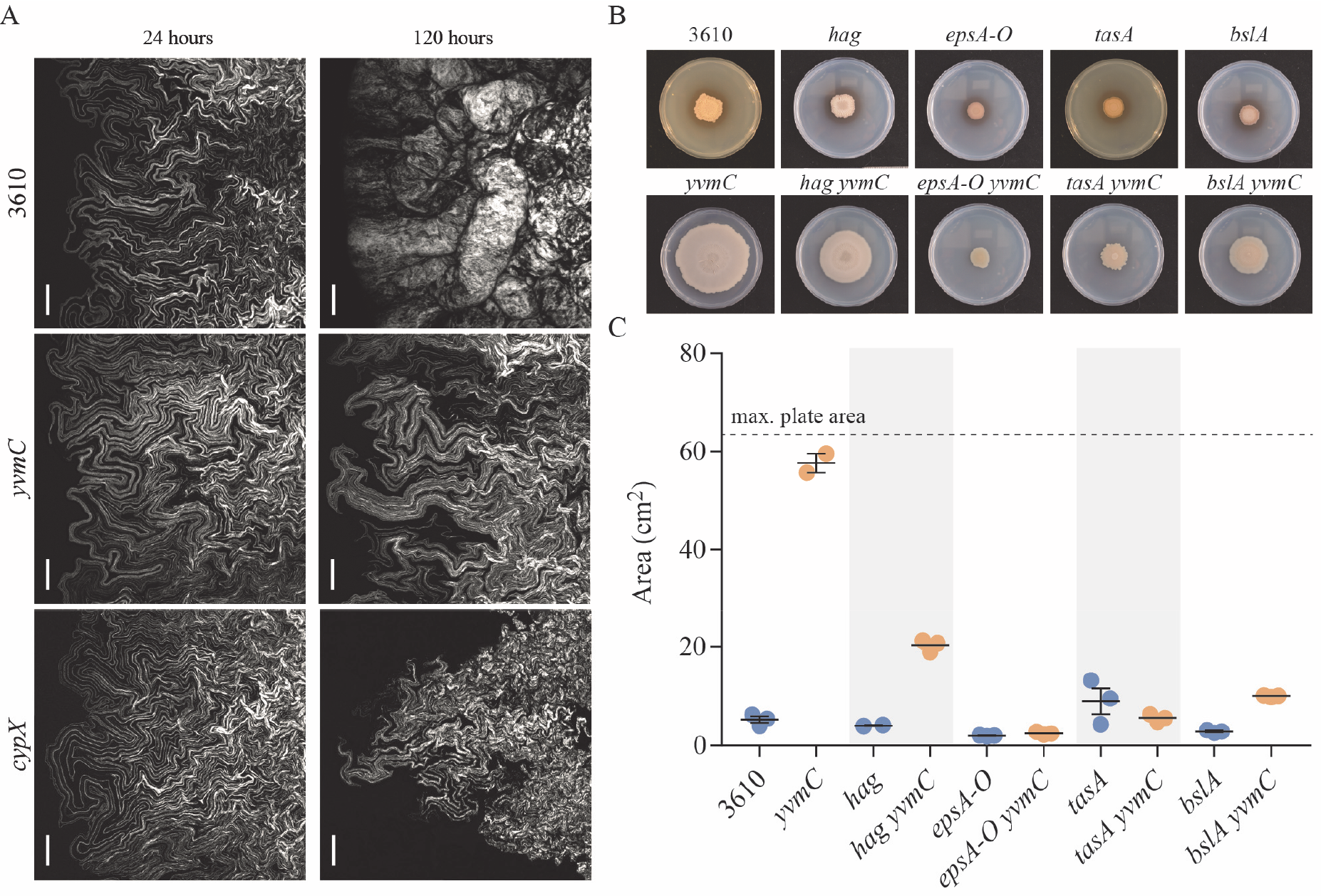
Biofilm expansion in the absence of pulcherriminic acid production. **(A)** Confocal microscopy of the biofilm edge at 24 and 120 hours for NCIB 3610 and the *cypX* (NRS5532) and *yvmC* (NRS5533) deletion strains. In each case the strains were mixed with an isogenic variant carrying a constitutively expressed copy of *gfp* (see methods and Table S1). This allowed detection of a fraction of the cells in the biomass by confocal microscopy. The images shown are projections of the acquired z-stacks and the scale bars represent 100 μm. In each case the outer edge of the biofilm is on the left hand side; **(B)** Biofilms formed by strains NCIB 3610, *hag* (DS1677), *epsA-O* (NRS2450), *tasA* (NRS5267), *bslA* (NRS2097), *yvmC* (NRS5533), *yvmC hag* (NRS5560), *yvmC epsA-O* (NRS6269), *yvmC tasA* (NRS6276) and *yvmC bslA* (NRS6281) were imaged after 120 hours; **(C)** The area occupied by biofilms formed by strains detailed in (B) on the 9 cm diameter petri dish was calculated and plotted. The bar represents the average of the biological repeats that are presented as individual points, the error bars are the standard error of the mean. The samples were grown at 30°C on MSgg agar containing 50 μM FeCl_3_.

Expansion of the *B. subtilis* biofilm across a surface has been shown to be driven by pressure exerted by the components in the biofilm matrix, rather than associated with flagellar-mediated motility (24). To determine whether the same processes drive radial expansion of the biofilm in the absence of pulcherriminic acid synthesis, mutations in *tasA, epsA-O, bslA* and *hag* (which encodes flagellin) were introduced into the *yvmC* mutant (Table S1). The footprint of the resultant biofilm was quantified after 120 hours incubation and the isogenic pulcherriminic acid positive strains were used as a reference for measuring biofilm expansion in the presence of pulcherrimin formation. We found that insertion of either the *epsA-O* or the *tasA* mutation to the *yvmC* mutant completely blocked expansion of the pulcherrimin deficient strain (Fig. 3B,C), with the *epsA-O* and *tasA* deletion strains occupying the same limited area of the surface whether or not pulcherriminic acid was made (Fig. 3B,C). However, for the *bslA* mutation, a small level of biofilm expansion was still attainable in the absence of BslA when pulcherrimin was not deposited into the agar (Fig 3B,C). Finally, the role of flagellar based motility in expansion was tested. Consistent with previous findings (24), the footprint of the *hag* biofilm was comparable to that of the parental NCIB 3610 biofilm (Fig. 3B,C). In contrast, while the area covered by the *yvmC hag* biofilm was significantly reduced by comparison with the *yvmC* mutant, it was greater than that occupied by the pulcherriminic acid positive *hag* deletion (Fig. 3B,C). The colonisation the pulcherrimin non-producing strains across a surface is therefore the consequence of a combination of flagellar based motility coupled with expansion driven by the biofilm matrix molecules.

### Cytoplasmic pulcherriminic acid does not trigger an arrest in biofilm expansion

We next wanted to elucidate whether cytoplasmic pulcherriminic acid or extracellular pulcherrimin was responsible for the observed biofilm growth arrest. For this purpose we needed to generate a strain where secretion of pulcherriminic acid was prevented. In *B. licheniformis*, the major transmembrane transporter of pulcherriminic acid is YvmA (30) and a homologue with 59% protein identity is encoded on the genome next to *cypX* and *yvmC* in *B. subtilis*. We therefore deleted *yvmA* and assessed pulcherriminic acid secretion by monitoring pigment production. Consistent with an involvement in pulcherriminic acid secretion, the agar under cells carrying the *yvmA* deletion was a very light brown, not the rusty brown observed for the wild-type strain or the clear agar of the *cypX* and *yvmC* deletion strains (Fig. S4). These findings are consistent with YvmA being the critical, but perhaps not sole, transporter of pulcherriminic acid to the extracellular environment. (The small amount of pulcherrimin in the agar is also consistent with limited cell lysis releasing low amounts of pulcherriminic acid into the external environment.) Having linked YvmA to release of pulcherriminic acid from the cell, we tested if the *yvmA* deletion strain formed a biofilm that was constrained in terms of the area it occupied or if the community kept expanding. The analysis determined that the biofilm formed by the *yvmA* mutant kept expanding, reaching levels similar to those measured for the *cypX* and *yvmC* strains (Fig. 2H and S2A). These results denote that the synthesis of intracellular pulcherriminic acid in itself is not sufficient to prevent expansion of the biofilm. Rather, it is the export of pulcherriminic acid to the environment and the subsequent formation of pulcherrimin that causes biofilm growth arrest.

### Iron limitation occurs at the edge of the wild type biofilm

As pulcherrimin is a complex between Fe^3+^ and pulcherriminic acid, one mechanism to explain the growth arrest of the wild-type biofilm is localised nutrient deprivation in the form of iron limitation. To test this hypothesis, we first constructed a mathematical model that incorporates the core processes of biofilm growth and pulcherriminic acid production. In the model, biofilm growth is proportional to the free iron concentration. Expansion of the biomass is driven by pressure resulting from cell division and the production of the extracellular matrix (31). Pulcherriminic acid is assumed to be produced and secreted by cells in the biofilm (see S text) and synthesis can be modulated to mimic the impact of the *yvmC* and *cypX* gene deletions in *B. subtilis*. Subsequent diffusion of pulcherriminic acid drives the chelation of iron in the growth medium and results in the formation of pulcherrimin (see supplemental text) (Fig. 4A). With parameters set to represent the wild-type strain, the output of the model manifests as an initial linear growth phase that is followed by a deceleration of biofilm expansion leading to a fixed biofilm size (Fig. 4B). When the level of pulcherriminic acid production is reduced below a critical level (to mimic either the *yvmC* or *cypX* mutant strains), the model again reveals a biphasic growth pattern. However, for the model mutant the initial linear growth phase is followed by seemingly unlimited exponential expansion (Fig. 4B). Therefore, this minimal mathematical model is capable of capturing the global growth characteristics that were measured experimentally for both the wild-type strain and those unable to synthesise pulcherriminic acid. The model output supports our hypothesis that arrest of biofilm expansion is mediated by a simple process that is influenced by just one molecule, pulcherriminic acid. In particular, the model predicts that pulcherriminic acid diffuses beyond the footprint of the biofilm and it is this that causes the arrest in biofilm expansion. A pulcherriminic acid “wave” is predicted to overtake the leading edge of the biofilm, culminating in a zone of iron limitation in the agar beyond the expanding biofilm edge (Fig. 4C).

**Figure 4:**
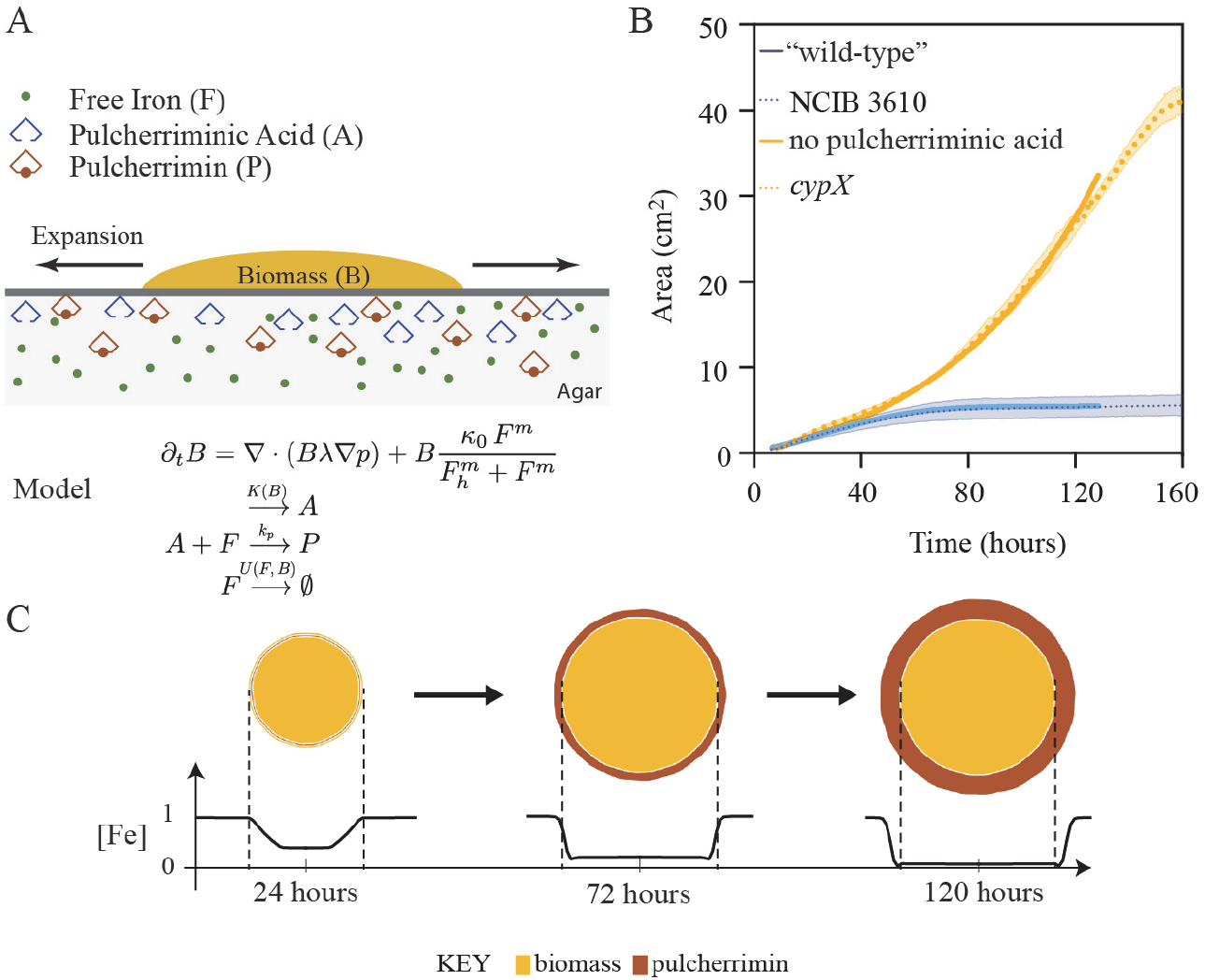
Mathematical model reveals iron limitation beyond the biofilm edge. **(A)** Schematic of the model set-up. In the model the biofilm biomass (*B*) grows on an agar substrate containing iron and expands radially at a rate proportional to the free iron in the media (*F*). Free iron is utilised for biomass production and maintenance at a rate *k_f_F.B*. Pulcherriminic acid (*A*) is produced and secreted by cells in the mature biofilm at a rate *k_b_B*. The pulcherriminic acid chelates iron with rate *k_p_A. F* resulting in the formation of pulcherrimin (*P*); **(B)** The area of the biofilm is plotted over time. Both the biological and mathematical data are shown. The dotted line is the average of 3 biological repeats and shadowed area is the standard error of the mean [the NCIB 3610 and *cypX* data are from Figure 2(H)]. The solid lines show the best fit of the model in the presence and absence of pulcherriminic acid production; **(C)** Model output showing biofilm expansion and production of pulcherrimin over time (*top view);* and level of free iron remaining in the agar (initial level normalised to unity). Three different time points are shown.

To test the ‘depletion wave’ hypothesis, we measured the level of available iron in the agar where the pulcherrimin was located after 48 and 72 hours (Fig. 5A). While the uncolonised agar retained an iron level of approximately 50 μM, the level of available iron in the agar where the pulcherrimin was deposited dropped to below the detection limit of the assay (< 1 μM) by 48 hours (Fig. 5A). In contrast, in an equivalent area of agar taken from under the biofilm formed by the *yvmC* and *cypX* deletion strains, approximately 15-20 μM iron remained, even after 72 hours incubation (Fig. 5A). To test whether the cells responded in distinct ways to these different iron levels we used a *PdhbA-lacZ* transcriptional reporter fusion. The *dhbA* gene encodes a protein needed for the biosynthesis of the siderophore bacillibactin which is synthesized under conditions of iron limitation (32) (33) (Fig. 5S). For the wild type strain evidence of activity from the *PdhbA-lacZ* transcriptional reporter could not be detected at early time points of biofilm formation (up to 48 hours) (Fig. 5B, 5C). However, around 72 hours, the time at which the cells cease expanding across the surface, a ring of cells with a deep blue colour developed near the periphery of the biofilm, which is indicative of *dhbA* transcription (Fig. 5D). This pattern of colouration was sustained and enhanced in the biofilm at later time points (Fig. 5E). In contrast, for both strains that do not form pulcherrimin, very little or no blue colour was visible in the biofilm even after 120 hours, indicating that transcription of the *PdhbA-lacZ* fusion was negligible (Fig. 5F, 5G). Thus, the wild-type biofilm experiences the iron deprivation particularly towards the edge of the biomass, whereas the pulcherrimin deficient strains are not exposed to the same degree of iron limitation. Collectively these findings show that while cell growth consumes iron from the agar, it is pulcherrimin production that results in the reduction of free iron in the biofilm vicinity to levels that cannot support continued expansion.

**Figure 5:**
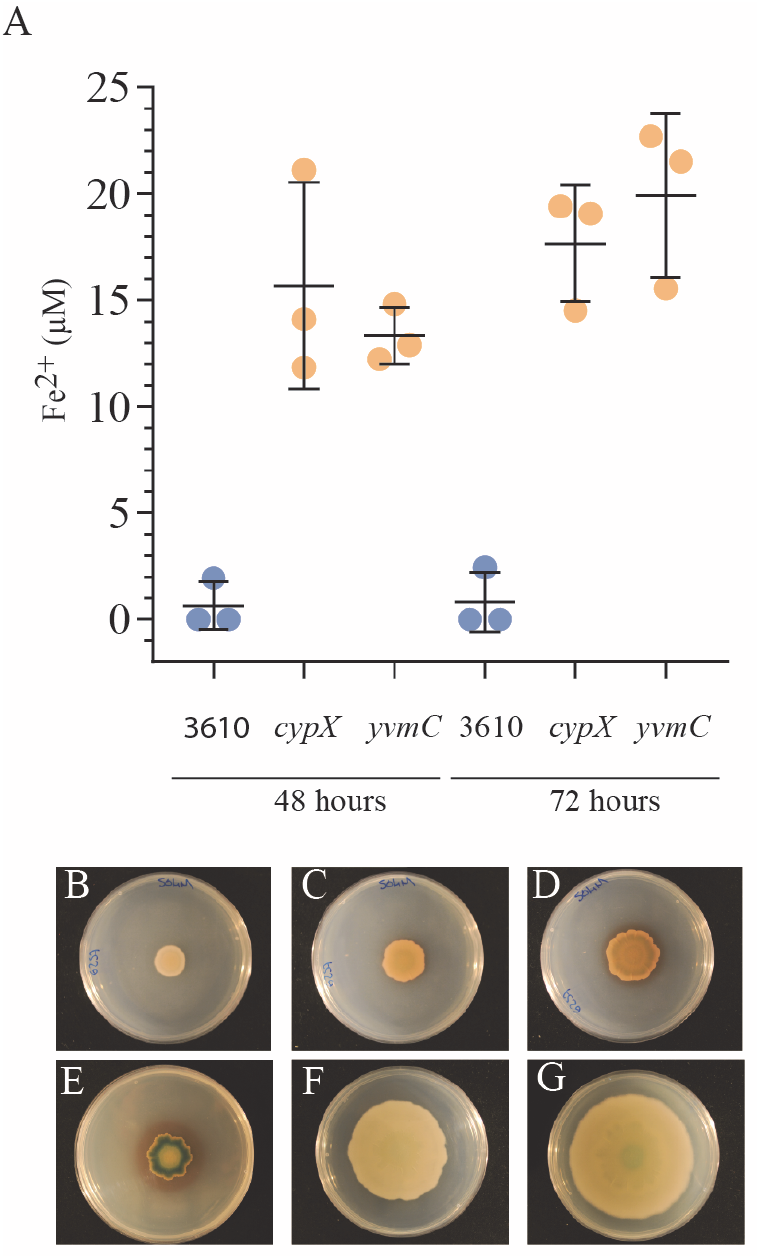
Detection of iron depletion is heterogeneous in the population. **(A)** The level of available ferrous and ferric iron in the agar was detected using a ferrozine assay. The bar represents the average of the biological repeats that are presented as individual points, the error bars are the standard deviation. Biofilms formed by NCIB 3610 and the *cypX* (NRS5532) and *yvmC* (NRS5533) deletion strains were grown at 30°C on MSgg agar containing 50 μM FeCl_3_ for the times indicated prior to analysis of the agar; **(B-G)** Images of strains containing the *PdbhA-lacZ* transcriptional reporter fusion after growth at 30°C on MSgg agar in 9 cm diameter petri dishes containing 120 μg ml^−1^ 5-bromo-4-chloro-3-indolyl-β-D-galactopyranoside (X-gal). **(B-E)** 3610 *PdbhA-lacZ* after 24, 48, 72 and 120 hours incubation respectively; **(F)** *cypXPdbhA-lacZ* at 120 hours incubation and; **(G)** *yvmC PdbhA-lacZ* at 120 hours incubation The image shown is representative of three biological repeats.

### Expansion of the biofilm is restricted at extremes of iron levels

Based on the data presented above we explored whether increasing the level of iron in the growth medium could overcome the arrest in biofilm expansion seen for the wild-type strain. We first tested this using the mathematical model. The output revealed that increasing the iron concentration modulated the biphasic growth profile leading to a longer initial linear phase (Fig. S6A) with the terminal area occupied by the biofilm increasing in proportion to the initial iron level (Fig. S6B). We note that the model predicted that increasing the initial level of iron in the environment constrains the pulcherrimin halo closer to the footprint of the biomass during expansion phase and at the terminal growth point (Fig. S6C). Following the trend of the mathematical data, our experimental data show that when the level of FeCl_3_ in the growth medium was raised from 50 μM to 500 μM the footprint of the biofilm formed by NCIB 3610 also increased (Fig. 6A, S6D). However, in contrast to the mathematical predication, no distinct halo of pulcherrimin protruded beyond the edge of the biofilm. Instead we noted that pulcherrimin was prevalent both within the biomass of the biofilm (which was dark pink in colour) and in the agar. The depletion of iron from the growth medium upon production of pulcherrimin was substantial. When we measured the level of available iron in the agar underneath the wild-type biofilm grown on a starting FeCl_3_ concentration of 500 μM, it was approximately 120 μM at 48 hours and dropped to below the detection limit of the assay by 72 hours (Fig. 6B). These data demonstrate that increasing the iron in the environment can overcome the impact of iron chelation by pulcherriminic acid.

**Figure 6:**
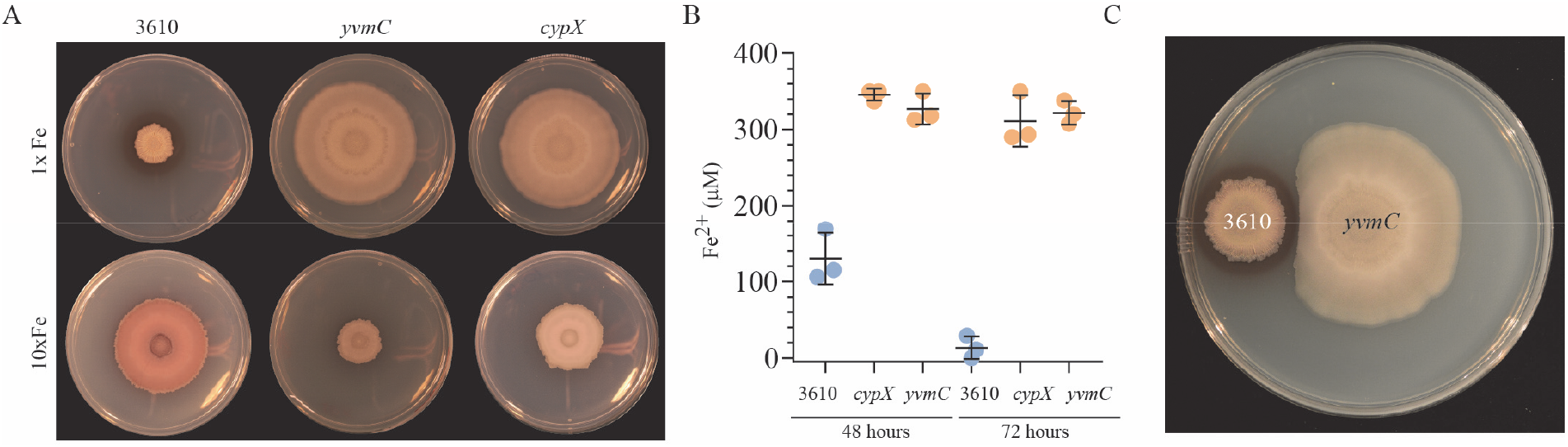
Pulcherriminic acid production modulates local iron levels. **(A)** NCIB 3610 and the *cypX* (NRS5532) and *yvmC* (NRS5533) deletion strains were grown at 30°C on MSgg agar containing either 50 μM FeCl_3_ (1x) or 500 μM FeCl_3_ (10x) for 120 hours prior to imaging; **(B)** The level of available ferrous and ferric iron in the agar was detected using a ferrozine assay. The bar represents the average of the biological repeats that are presented as individual points, the error bars are the standard deviation. The starting concentration in the agar was 500 μM FeCl_3_ and the maximum detection limit in these conditions was 350 μM; **(C)** Biofilms of NCIB 3610 and *yvmC* (NRS5533) were inoculated onto a 15 cm diameter MSgg (1.5% w/v agar) plate and incubated at 30°C for 120 hours prior to imaging.

In contrast to the rapid depletion of iron in the agar in the vicinity of the wild-type strain, we found that the agar under the biomass of *yvmC* and *cypX* deletion strains retained approximately 300-350 μM iron, even after 72 hours incubation (Fig. 6B). Additionally, and again contrary to that of the wild-type strain, analysis of biofilm formation by the *yvmC* and *cypX* deletion strains revealed that the 10-fold increase in the FeCl_3_ level resulted in a reduction of the footprint area occupied after 120 hours (Fig. 6A, S6D). Based on these findings, we hypothesized that the negative impact on footprint area of increasing iron concentration in the medium could be a consequence of the level of iron in the growth medium being toxic for the *yvmC* and *cypX* deletion strains. However, analysis of cells grown in planktonic conditions revealed that the growth rates of the *yvmC* mutant was broadly similar to that of the wild-type (Fig. S6E). Thus, the lack of biofilm expansion in the presence of 500 μM FeCl_3_ for strains that do not produce pulcherriminic acid is not a consequence of toxicity. Therefore we conclude that when the level of iron is above a threshold the cells are prevented from triggering expansion of the biofilm. When these findings are coupled with the self-restriction of growth of the wild type strain when cultured on 50 μM FeCl_3_, they indicate that if the level of available iron in the local environment is either too low or too high expansion of the biofilm is restricted.

### Protection of a niche through pulcherrimin formation

As pulcherrimin restricts growth through local depletion of iron, we hypothesised that production of pulcherriminic acid might be advantageous to protect a local niche. We therefore explored if the pulcherrimin deposit produced by the wild-type biofilm could restrict the growth of neighbouring populations. When the wild-type and *yvmC* mutant strain were inoculated onto the same plate, we found that the otherwise unlimited growth of the *yvmC* biofilm was restricted just in the proximity of the pulcherrimin in the agar (Fig. 6C). In the analogous experiment where the *yvmC* mutant was grown adjacent to another biofilm formed by the *yvmC* mutant there was no zone of growth inhibition (Fig. S6F), with cells growing abutted to each other. Thus, pulcherrimin-forming populations can protect their niche from colonisation by neighbouring bacteria.

## Discussion

By coupling mathematical modelling and microscopy with bacterial genetics, we have revealed that growth arrest is a distinct stage of *B. subtilis* biofilm development that is not simply a consequence of spore formation. Our data show that a fraction of the cells remain metabolically active even though expansion of the biofilm has stopped. We identify that production of pulcherriminic acid underpins growth arrest of the biofilm. However, its synthesis is not sufficient; it is the chelation of Fe^3+^ through the formation of pulcherrimin surrounding the population on the biofilm that is needed to self-restrict biofilm expansion. Self-restriction of growth by a molecule produced by the same population of bacteria is counterintuitive from an evolutionary perspective and this led us to explore possible advantages associated with formation of pulcherrimin. Our analyses revealed beneficial effects for the pulcherrimin-producing bacterium as the pulcherrimin extending outside the biomass of the biofilm can prevent colonisation by neighbouring communities. We also uncovered that pulcherrimin formation confers an advantage through modulation of the external conditions in high iron environments.

### Growth in the biofilm is controlled by iron levels

Iron is a micronutrient that is essential to support bacterial growth and survival, but while it is relatively abundant in the soil it is largely inaccessible (34). The low accessibility of iron within the soil stimulates competition between plants and microbes for its acquisition. Bacteria have evolved many mechanisms to capture and sequester iron from the environment including iron uptake systems (35), siderophore production (32) and sorption methods (36). Here we have uncovered the contribution that iron chelation, mediated by pulcherrimin formation, has on the physiology of the *B. subtilis* biofilm. Indeed, we have shown that chelation is the dominant cause of free iron depletion from the medium - not utilisation for growth itself. At low iron levels pulcherriminic acid diffuses beyond the expanding biofilm leading edge, chelating Fe^3+^ and self-restricting expansion of the biomass. These findings are consistent with data linking *B. subtilis* biofilm formation to iron availability (37, 38). Consistent with iron levels being the dominant factor influencing biofilm expansion, both mathematical and experimental data show that increasing the iron concentration in the growth medium overrides this growth inhibition. When iron is plentiful, the profile of pulcherrimin deposition alters and a halo surrounding the cells is not apparent. This means that the leading edge of the biofilm is not exposed to iron limitation and continues to expand for longer. The molecular mechanisms controlling the transition of *B. subtilis* to a non-expanding phase when iron levels are too low needs to be elucidated. Moreover, whether there is a feedback loop between the concentration of iron in the environment and pulcherriminic acid production or if pulcherriminic acid biosynthesis is independent of the level of iron needs further investigation. The possibility of post-transcriptional regulation of pulcherriminic acid production is supported by the identification of an extended 5’ untranslated region upstream of the *yvmC* coding region (39). Such regions are typical of riboswitches that can bind a wide array of metabolites and metals and control gene transcription and translation efficiency for example (40, 41).

### Sequestration of iron in biofilms

The ability to modulate iron levels in the context of biofilm formation appears to be a theme emerging in the literature. While in this study we have uncovered the involvement of iron chelation by pulcherrimic acid in growth arrest, the accumulation of ferric and ferrous ions in the matrix of *B. subtilis* has been linked with protection from erosion and toxicity (42). The sequestration of iron within the extracellular matrix is not unique to *B. subtilis* as in biofilms formed by the Gram-negative bacterium *Pseudomonas aeruginosa* the Psl polysaccharide fibres of the matrix bind and sequester both ferric and ferrous ions. This is not a passive process as high levels of iron in the environment trigger Psl synthesis and therefore promote accumulation of iron in the local environment of the bacterial population (36). Similarly a polysaccharide produced by *Klebsiella oxytoca* biofilms has also been shown to bind iron (43) and electron dense iron-enriched deposits accumulate in the fibrous matrix of *Enterococcus faecalis* biofilms (44). In *E. faecalis* the deposits promote extracellular electron transport by acting as an electron sink and thus support growth in the biofilm through altered metabolism. Therefore the purpose of controlling iron levels in the microenvironment of the biofilm is varied and is likely to be specific for different species.

### Pulcherriminic acid mediated chelation of iron

Pulcherrimin has been observed to be made by both eukaryotic and prokaryotic microorganisms (27, 45). It is, however, only more latterly that the biosynthetic pathways involved have been elucidated (46, 47). Our comparative genomic analysis revealed that the genes needed for pulcherriminic acid synthesis and secretion (*yvmC*, *cypX* and *yvmA*) were found in the genome or on plasmids of species beyond *B. subtilis*. It should be noted that the pulcherrimin biosynthetic cluster is not found in all isolates of each species (including *B. subtilis*), but those bacteria that contain the genes include isolates of *Bacillus cereus, Bacillus thuringenesis* and *Staphylococcus epidermidis* species (Fig 7A, B). Differences in gene synteny from that of the *B. subtilis* genome were apparent with the regulator, *pchR*, not always being present in the genome and additionally the organisation of the remaining genes was varied (Fig. 7B). However, one consistent feature in the gene organisation pattern was that the *yvmC* coding region always preceded that of *cypX*. The patchy distribution of the genes both within a single species and between closely related bacteria suggests loss of the genes overtime, rather than reoccurring acquisition. This pattern of gene loss and distribution is consistent with the profile observed in yeast (46). However, given the knowledge that at least two routes have evolved for pulcherriminic acid production (46, 47), we cannot exclude the possibility that other species also synthesize pulcherriminic acid using an as yet unidentified pathway.

**Figure 7:**
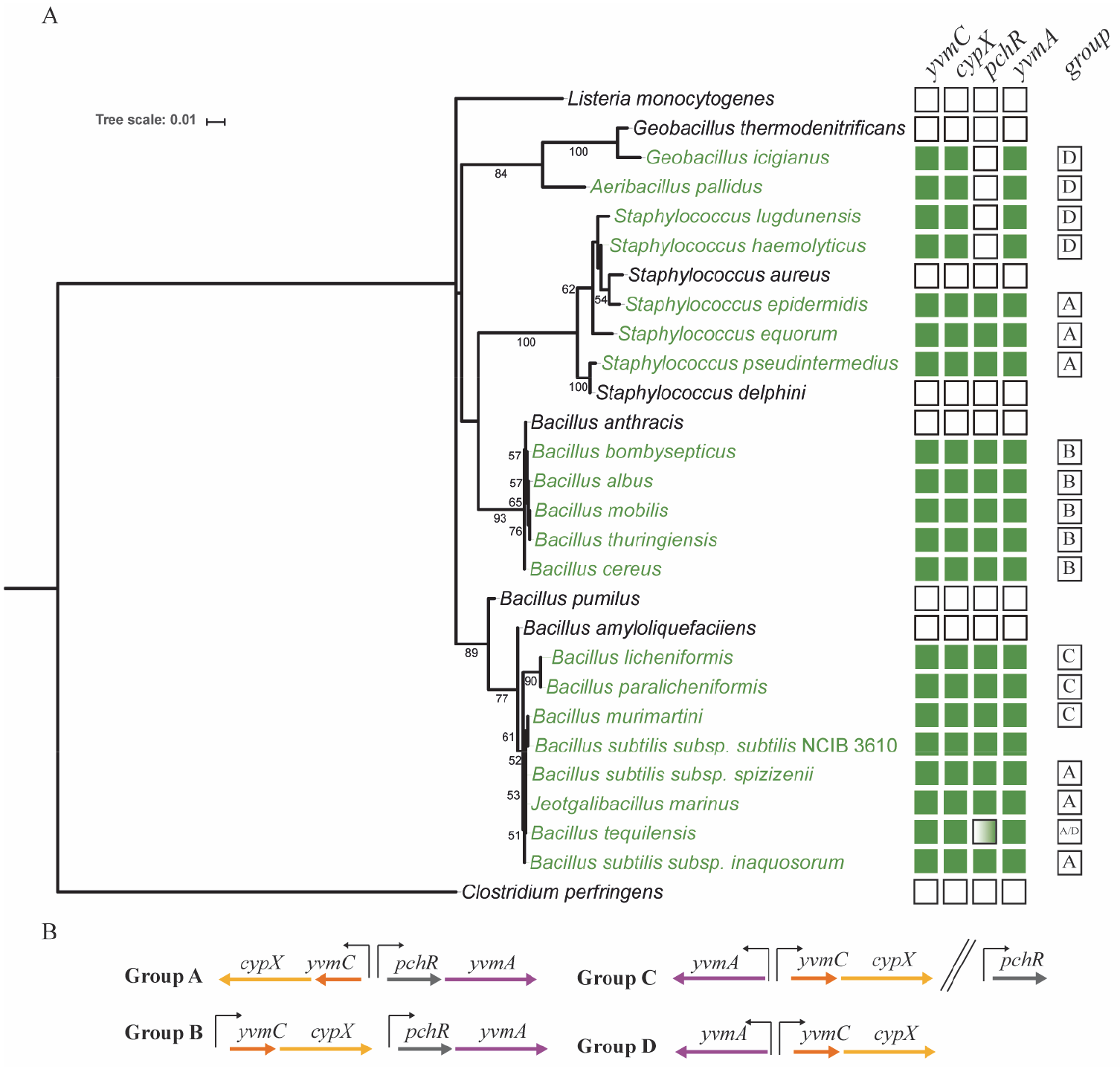
Comparative genomic analysis of the pulcherriminic acid biosynthetic cluster. **(A)** Phylogenetic tree showing species that contain genes within the pulcherriminic acid biosynthetic cluster. The numbers adjacent to branches indicate the bootstrap values for 350 replicates. Species with members that contain the cluster are shown in green and those without the genes in black. The boxes on the right hand side show the genes that were identified in the genomes, green is positive and white is negative. The dual green/white box indicates variability in the presence of the gene throughout the genomes. The box containing the letters A-D indicate the genomic organisation of the genes in the cluster; **(B)** The organisation of the pulcherriminic acid biosynthetic genes as identified through comparative genome analysis. The black upright arrows represent promoters and the two parallel slanted lines indicate a gap in the genome between the genes.

In yeast strains, such as *Metschnikowia pulcherrima*, pulcherrimin formation has been linked with antagonism against other species (48). In *B. subtilis*, the function of pulcherrimin was entirely unknown. We have identified that pulcherriminic acid allows *B. subtilis* to alter the level of freely available iron in its vicinity, over (at least) two-orders of magnitude, as an initial concentration of 500 μM FeCl_3_ in the growth medium can be entirely sequestered within 72 hours (Fig. 6B). This remarkable ability to manipulate the external environmental conditions has corresponding benefits. For example, the pulcherrimin halo of the wild-type strain prevents invasion of its niche by bacteria found in the surrounding environment (Fig. 6C). This niche protection mechanism is likely to be effective against any microorganism that depends on iron for growth. Consistent with this, pulcherrimin production has been associated with biocontrol processes (49). However, this long held interpretation of how pulcherrimin functions has been challenged by the recent classification of pulcherriminic acid as a siderophore in the yeast *Kluyveromyces lactis* (46). In *K. lactis* a membrane bound protein encoded by the gene PUL3 is linked with utilisation of pulcherrimin via uptake. This opens up the possibility that pulcherrimin producers across both bacterial and eukaryotic microbes are exploited by neighbouring “cheater” cells. In this context, the non-producing cells could make use of the chelated iron but would not contribute to its deposition (46). In nature, the routes used for iron acquisition are highly diverse and competition between ecologically distant species is common. For instance, insect herbivores are able to hijack plant iron acquisition systems (50) and the soil bacterium *Streptomyces venezuelae* produces a volatile compound (trimethylamine) that enhances colonisation by the bacterium through modulating iron availability (51, 52). Therefore, while there is no evidence of *B. subtilis* being able to use pulcherriminic acid as a siderophore, the intriguing question of how pulcherrimin is accessed and utilised in the complex soil environment remains open to multiple possibilities.

## Materials and Methods

### General growth conditions and strain construction

The *Bacillus subtilis* strains used and constructed in this study are detailed in Table S1. *B. subtilis* 168 derivatives were obtained by transformation of competent cells with plasmids using standard protocols (53). SPP1 phage transductions were used to introduce DNA into *B. subtilis* strains (54). *Escherichia coli* strain MC1061 was used for the construction and maintenance of plasmids. *E. coli* and *B. subtilis* strains were routinely grown in Lysogeny-Broth (LB) medium (10 g NaCl, 5 g yeast extract, and 10 g tryptone per litre) at 37°C for 16 hours. When appropriate, antibiotics were used at the following concentrations: ampicillin 100 μg ml^−1^, erythromycin 1 μg ml^−1^, kanamycin 25 μg ml^−1^, neomycin 8 μg ml^−1^ and spectinomycin 100 μg ml^−1^.

### Plasmid construction

All primers used in this study are presented in Table S2. Genetic complementation of Δ*yvmC* and Δ*cypX* was achieved by PCR amplification of the *yvmC-cypX* region, including 500 base pairs upstream of the *yvmC* coding region (using primers NSW2461 and NSW2456), from NCIB 3610. Fragment was digested using the EcoRI/SphI restriction enzymes and the purified fragment was ligated into pDR111. The resulting plasmid pNW1722 was introduced to *B. subtilis* strain 168 at the *amyE* locus by virtue of the spectinomycin resistance cassette. The complementation construct was subsequently transferred to the NCIB 3610 derived strains Δ*yvmC* (NRS5533) and Δ*cypX*(NRS5532) using SPP1 phage transduction.

Activity from the *dhbA* promoter was monitored using production of β-galactosidase as the reporter through construction of a P*dhbA-lacZ* transcriptional fusion. The DNA carrying the promoter region was amplified by PCR using primers NSW2477 and NSW2478 with genomic DNA extracted from NCIB 3610 being used as the template. The PCR fragment was digested using the EcoRI and BamHI restriction enzymes and ligated into pDG1728. The resulting plasmid pNW1725 was introduced into the genome of 168 and subsequently introduced into the genomes of NCIB 3610, Δ*yvmC* (NRS5533) and Δ*cypX*(NRS5532) using SPP1 phage transduction.

### Biofilm formation

*B. subtilis* strains were grown on MSgg medium (5 mM potassium phosphate and 100 mM MOPS at pH 7.0 supplemented with 2 mM MgCl_2_, 700 μM CaCl_2_, 50 μM MnCl_2_, 1 μM ZnCl_2_, 2 μM thiamine, 0.5% glycerol, 0.5% glutamate) (9) solidified with 1.5% select agar (Invitrogen) at 30 °C at the indicated time points. FeCl_3_ was added at the indicated concentration and where not specified was used at 50 μM. When appropriate 5-bromo-4-chloro-3-indolyl ß-D-galactopyranoside (X-Gal) was added at 120 μg ml^−1^. To set up a biofilm, a 3 ml aliquot of LB medium was inoculated with an individual colony taken from an overnight plate and grown at 37°C to an OD_600_ of 1. Unless otherwise stated, 5 μl of the culture was placed onto an MSgg plate which was incubated at 30°C for morphology and hydrophobicity studies. Images of colony biofilms were recorded using a Nikon D3200 digital camera mounted on a Kaiser RS3XA copy stand or using a Leica MZ16FA stereomicroscope.

### Footprint area measurements

Images were taken using either a 5-megapixel USB autofocus camera module CAM8200-U controlled with python-bash software developed specifically or using a BioRad GelDoc XR+. Images were analysed with FIJI (55) using standards thresholding methods (56) to extract the biofilm footprint area occupied.

### Sporulation assay

For heat resistant spore quantification, colony biofilms were grown for 24, 48, 72, 96 or 120 hours at 30°C. The complete biofilm was harvested and suspended in 1 ml of saline solution. The biomass was disrupted by passage through a 23 X 1 needle 8 times and subsequently subjected to mild sonication (20% amplitude, 1 second on, 1 second off for 5 seconds total) to liberate bacterial cells from the matrix. To kill vegetative cells, the samples were incubated for 20 min at 80°C. To determine the percentage spores, serial dilutions were plated before and after the 80°C incubation on LB agar supplemented with 25 μg ml^−1^ kanamycin. The percentage of spores was established by colony forming unit counting and results are presented as the percentage of colony forming units obtained after incubation of the samples for 20 min at 80°C divided by the number of colony forming units obtained before the heat inactivation.

### Biofilm hydrophobicity imaging

Biofilm hydrophobicity was determined by placing a 5 μl droplet of water on the upper surface of biofilms that had been grown for 48 hours at 30°C (17). The water droplet was allowed to equilibrate for 5 minutes prior to imaging using a ThetaLite TL100 optical tensiometer (Biolin Scientific).

### Ferrozine assay

The level of Fe^2+^ present in the agar was measured as follows. A “punch” of agar was taken using the reverse side of a 200 μl pipette tip and the extracted agar was expelled into a well of a 96-well plate. The absorbance at 562 nm was measured and recorded to provide the baseline. The assay was started by addition of 63 μl of the master mix that comprised 46.7 μl of 1 M potassium acetate (pH 5.5 with acetic acid), 3.3 μl 1 M ascorbic acid to reduce available Fe^3+^ to Fe^2+^ and 13 μl ferrozine (stock 50 mg/ml in 1 M potassium acetate). The samples were incubated at 37°C for 1 hour before reading absorbance at 562 nm using a plate reader. A standard curve for the assay was generating using agar punches extracted from MSgg agar plates prepared with defined concentrations of FeCl_3_ that ranged from 0 to 400 μM. The maximum value of Fe^2+^ that could be measured using this method was 350 μM, therefore samples that presented as above this value were given a value of 350 μM as this represents the upper limit of the assay. The data presented are an average of three technical samples for each condition.

### Confocal Microscopy

4 ml of MSgg medium supplemented with 1.5% (w/v) agar was placed into a 35 mm diameter petri dish and dried for 1 hour in a laminar flow hood. NCIB 3610, Δ*cypX* and Δ*yvmC* strains and their respective constitutive GFP-producing daughter strains (Table S1) were grown separately in LB medium to an OD_600_ of 1. The cultures were then mixed to produce a suspension of cells where 80% were non-fluorescent and 20% GFP expressing. 1 μl of this mixed culture solution was spotted into the centre of the petri dish and was incubated at 30°C for the indicated time period.

A Leica SP8 upright confocal was used to image the edge of the biofilm using a 10x 0.3 N.A. air objective and a heated chamber that was pre-warmed to 30°C. A cling film tent was draped from around the objective and tucked loosely under the stage to eliminate air-flow across the plate and minimise dehydration (and therefore shrinkage) of the agar. An additional 35 mm diameter petri dish was filled with water and placed next to the biofilm plate to increase the humidity inside the tent. An argon-ion laser was used to excite the GFP at 488 nm and 2% power. Z-stacks capturing the full height of the biofilm border were specified based on the presence of GFP-containing cells and planes of 1024x1024 pixels were acquired quickly using a resonant mirror, averaging 16 scans per line. Images were imported into an OMERO (57) server and figures were prepared using OMERO.figure (http://figure.openmicroscopy.org/).

### Bioinformatics Analysis

Genome sequences of 44046 strains, representing 8244 species, were downloaded from Ensembl Bacteria release 40 (58), and supplemented with the sequence of *Staphyloccocus delphini* 8086 (PRJEA8701; ENA Release 127). Nucleotide sequences of the genes comprising the pulcherriminic acid biosynthetic cluster were obtained from *Bacillus subtilis* BS49 (GCA_000953615.1; ENA Release 127) and corresponded to the following coordinates on the genome: *cypX* (negative strand) 3638633-3639850; *yvmA* (positive strand) 3641568-3642779; *yvmB* (also known as *pchR*) (positive strand) 3641038-3641547; and *yvmC* (negative strand) 3639866-3640612. The nucleotide sequenced were translated to peptide sequences using the transeq program from the EMBOSS package version 6.5.7.0 (59). A similarity search of the database was carried out for each gene using the tblastn program of NCBI BLAST+ 2.5.0 (60) using an e-value cut-off of 0.05. The resulting alignments were parsed using a custom biopython (61) script to identify strains with sequences where >50% of the length of the query sequence was represented. The orientation and organisation of the genes within the cluster were also compared in isolates carrying sequences with similarity to all four members of the cluster to identify isolates carrying the sequences on the same contig sequence, and the relative orientation of these. Our analysis revealed that the *pchR* gene, which encodes a transcription regulator, was not always found in the same genomic location (Group C) as the other genes in the biosynthetic cluster and in some cases was missing from the genome (Group D) (Fig. 7B). Therefore we deemed *yvmC, cypX* and *yvmA* as the core cluster and included strains where *pchR* was absent in our analysis (Fig. 7A). The source of each sequence (chromosomal or plasmid DNA) was determined from annotations in the database entries, where appropriate annotations were provided. To resolve non-specific taxonomic assignments in the databases, 16S rRNA gene sequences from all species were identified in the genome sequences by BLASTN searches using the *B. subtilis* BS49 16S rRNA as a query, and the resulting sequences classified using ExBioCloud.net (62) (May 2017 database release) (Table S3). To construct the phylogenetic tree the 16S rRNA sequences were aligned by MAFFT v7.407 (63) using the L-INS-i algorithm. A maximum likelihood tree was generated using RAxML 8.2.12 (64) using the GTRGAMMA model and rapid bootstrapping analysis with automatic frequency-based criteria. *Clostridium perfringens* was included as an out group in the analysis and used to root the tree.

## Acknowledgments

Work in the NSW, CEM, and FAD groups is supported by the Biotechnology and Biological Sciences Research Council [BB/P001335/1; BB/R012415/1]. MK is supported by a Biotechnology and Biological Sciences Research Council studentship [BB/M010996/1]. We would like to acknowledge the Dundee Imaging Facility, Dundee, which is supported by the ’Wellcome Trust Technology Platform’ award [097945/B/11/Z] and the ’MRC Next Generation Optical Microscopy’ award [MR/K015869/1]. We thank Prof. J. Helmann for helpful discussions.

## Author contributions

Conceived the project – SA, NSW

Performed experiments – SA, MP, MK, DM

Conducted ferrozine assays- MP

Conducted bioinformatics analysis – JA, MK

Analysed data – NSW, FAD, SA, MP, MK, DM. CEM

Wrote model – FAD, DM

Conducted model simulations -DM

Wrote paper – SA, FAD, CEM, NSW

## Supplemental text

### Mathematical Framework

The model builds on that developed in (23) and considers the interaction of the following basic processes (see Fig 4).

- Biomass production (cell division and matrix production) within the biofilm causes internal pressure that drives outward spreading
- The rate of biomass production is limited (only) by the available iron concentration
- The biomass produces pulcherriminic acid, which diffuses into the growth medium
- Pulcherriminic acid chelates free iron forming the compound pulcherrimin

### Biomass Dynamics

The biomass density (*B*) must obey the follow continuity equation (conservation of mass within any given area implies that biomass created must equal the next flux of material into the region plus the net production of biomass within that region): *∂_t_B* = −∇ · *J* + *G*, where *J* and *G* are the flux and production rate of *B*, respectively. The flux is generated implicitly by growth; cell division, the production of extracellular matrix and consequent water absorption induces a pressure that expands the biomass. We therefore assume that the flux can be represented by the expression *B**ν*** with the velocity ***ν***, given by ***ν*** = −*λ*∇*p* where ∇*p* is the gradient of the pressure due to growth and *λ* is related to the mechanical properties of the material. This relationship is referred to as Darcy’s Law in certain contexts (see e.g. (65)).

The biomass production rate *G* is assumed to depend only on the availability of iron (i.e. iron is considered to be the growth-limiting substrate within the medium). In the absence of additional information, we initially assume biomass production follows a simple saturating response to iron availability. Moreover, the rate of production of biomass must depend on the presence of biomass and we assume this relationship to be linear in the first instance. Therefore we write

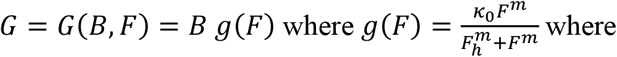

*κ*_0_ the maximal growth rate, *F_h_* represents the free iron level that corresponds to half maximal growth rate and *m* characterises the growth response. From the above, we can derive the biomass equation

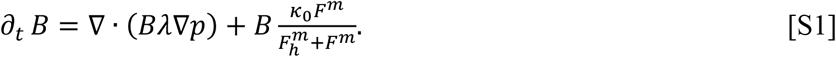

### Iron Dynamics

(i) Pulcherriminic acid is produced by the biomass. (ii) On diffusion into the agar, this acid irreversibly chelates iron to form pulcherrimin (see Fig. 4) (iii) Production and maintenance of biomass utilises iron:

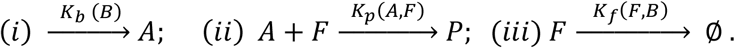

We assume simple relationships that follow the law of mass action: *K_b_* (*B*) = *k_b_B*, *K_p_* (*A, F*) = *k_p_ A. F* and *K_f_*(*F,B*) = *k_f_B* for acid production, pulcherrimin formation and iron utilisation, respectively with *k_b_,k_p_* and *k_f_* positive constants. Taking into account diffusion of the reactants in the agar (D_A_, D_P_, D_F_), we can model the above reactions with the following system of partial differential equations:

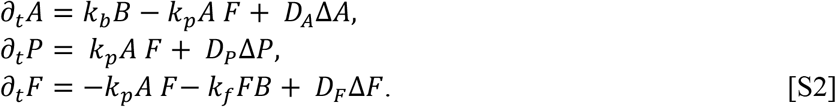

### The Moving Boundary Problem

Spatial and temporal scales ensure that it is reasonable to assume that the biofilm can be modelled as planar and within the expanding outer edge, *B* ≡ *B_S_* for some constant *B_S_*. Beyond the edge of the biomass, *B* ≡ 0 and hence the growth pressure *p* ≡ 0. Scaling also ensures that it is reasonable to assume there are no gradients in the *z* –direction. Finally, growth experiments demonstrate a high degree of radial symmetry. Therefore the model can be reduce further to a 1dimensional spatial domain where the space variable *r* represents radial distance from the centre of the expanding biomass. Eqns [S1] and [S2] become

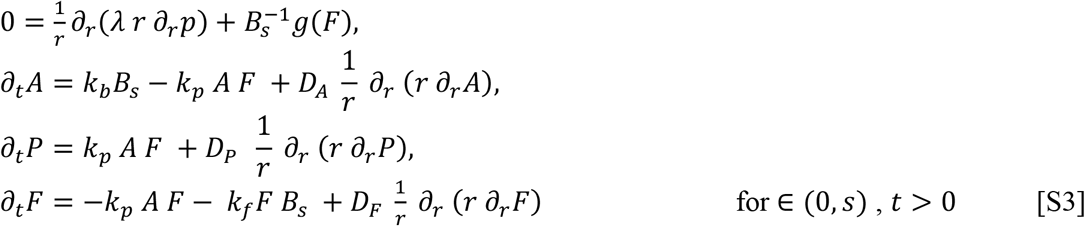

and

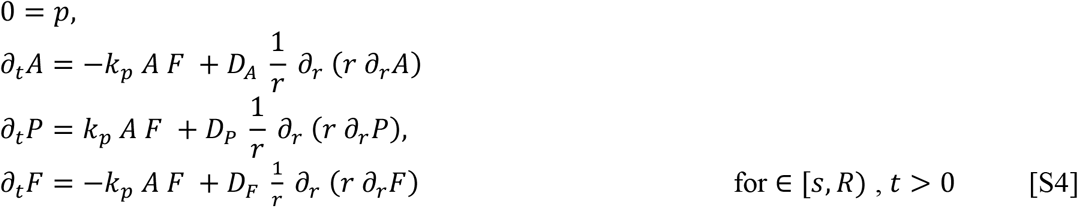

where *S* = *S*(*t*) marks the edge of the biofilm footprint and *R* is the radius of the Petri dish. Appropriate conditions at the moving boundary are

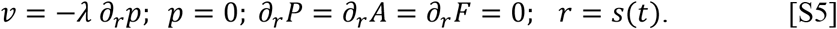

Initial data corresponding to a localised inoculum are:

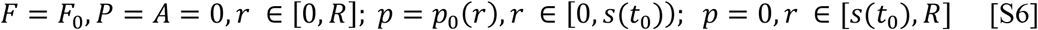

### Model Solution

The model was solved by first non-dimensionalising the dependent and independent variables. The following non-dimensionalisation was performed

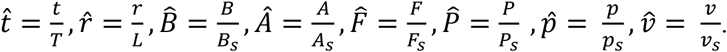

Setting 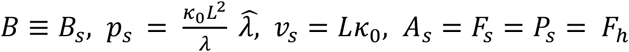 where *F_h_* is the level of free iron that results in the growth rate being half its maximal value results in a new set of grouped (non-dimensionalised) parameters given by

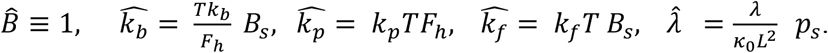

Our experiments suggested an appropriate time and length scale to be *T* = 250 mins and *L* = 1 mm capturing the observed initial maximal growth rate. Moreover, a reasonable estimation for the diffusion of pulcherriminic acid within media considered here is *D_A_* ~ 10^−2^mm^2^min^−1^. We therefore set 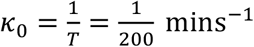 and 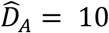. As a region of depleted iron is maintained within the footprint of the biofilm long after expansion has stopped, we make the reasonable assumption that the diffusion of free iron and pulcherrimin in the agar to be orders of magnitude smaller than that for pulcherriminic acid. Note that in the non-dimensional setting the first equation [S3] becomes

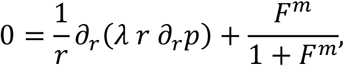

where we have dropped the hats for ease of exposition. Table S4 summarises the variables and parameters and the non-dimensional value used in the simulations (again dropping hats).

With non-dimensional variables and parameters replacing their counterparts in [S3-6] the system was solved using a finite difference method in space and time in an iterative manner to account for the moving boundary. At each step, the edge of the biomass was extended by a small amount governed by its velocity as computed by Eqn. [S5]. The well-known MATLAB solver pdepe (see e.g. https://uk.mathworks.com) was iteratively employed to implement a finite difference method of lines approach to produce solutions to each new (fixed) boundary value problem.

Scheme:

1. Set *t* = *t_0_* and set *S*(*t*) = 1. Choose a small number 0 < *δ* ≪ 1.
2. Prescribe the initial data [S6] with *p* = *p*_0_(*r*) > 0 and *F* = *F*_0_ >0 constant.
3. Compute *ν* = −*λ ∂_r_p* at *r* = *S*(*t*).
4. Set *S*(*t* + *δ t*):= *S*(*t*) + *νδt*.
5. Solve Eqns. [S3, S4, S5] for *p*, *P*, *A*, *F* with (*r*,*t*) ∈ [0, *R*] × (0, *t* + *δ t*].
6. Set *t* = *t* + *δ t* and return to 3.

**Table S1.**
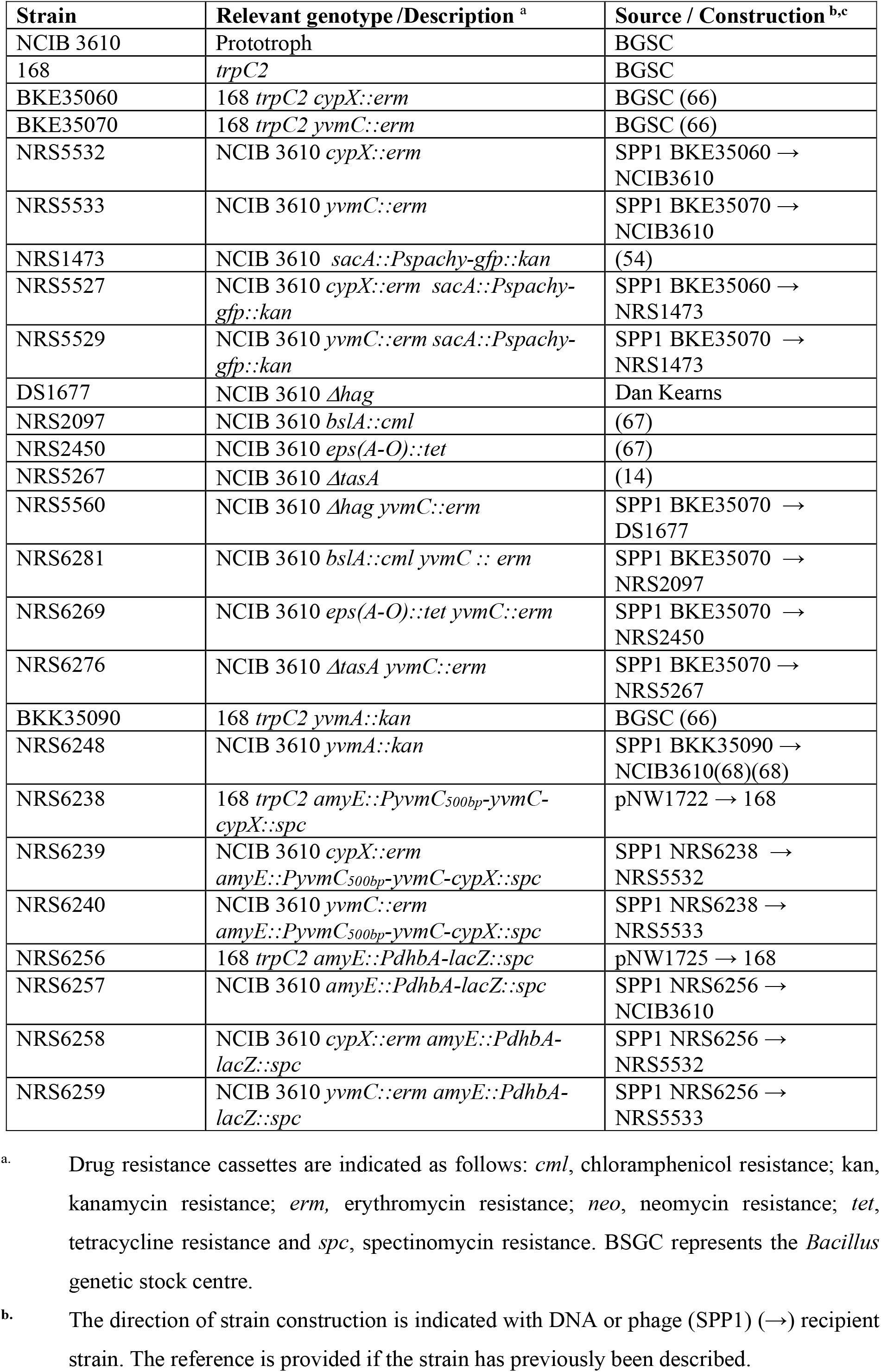
Full list of *B. subtilis* strains used in this study.

**Table S2:**
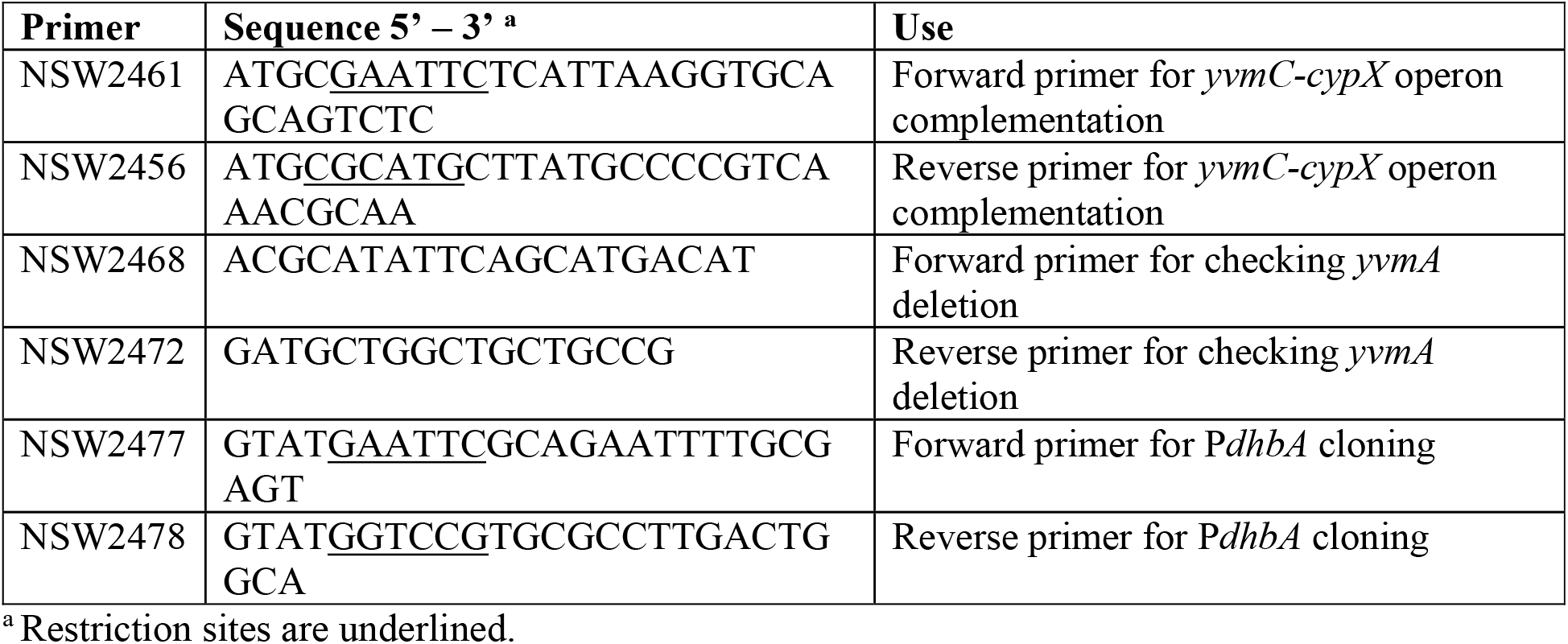
Oligonucleotide primers used in this study.

**Table S3:** Details of strains used for phylogenetic tree construction

| Species/strain identified by 16S analysis | Accession number | Database classification | Source |
| --- | --- | --- | --- |
| *Bacillus subtilis* subsp. *subtilis* NCIB 3610 | CP020102.1 | *Bacillus subtilis* subsp. *subtilis* NCIB 3610 | ENA Release 137 |
| *Aeribacillus pallidus* | GCA_001624605.1 | *Geobacillus* sp 8 | Ensembl Genomes release 40 |
| *Bacillus albus* | GCA_000238655.1 | *Bacillus* sp 7 6 55cfaa ct2 | Ensembl Genomes release 40 |
| *Bacillus amyloliquefaciens* | GCA_000696285.1 | *Bacillus amyloliquefaciens* | Ensembl Genomes release 40 |
| *Bacillus anthracis* | GCA_000725325.1 | *Bacillus anthracis* | Ensembl Genomes release 40 |
| *Bacillus bombysepticus* | GCA_000831065.1 | *Bacillus bombysepticus* | Ensembl Genomes release 40 |
| *Bacillus cereus* | GCA_000160915.1 | *Bacillus cereus* | Ensembl Genomes release 40 |
| *Bacillus licheniformis* | GCA_000952085.1 | *Bacillus licheniformis* | Ensembl Genomes release 40 |
| *Bacillus mobilis* | GCA_001429465.1 | *Bacillus* sp root11 | Ensembl Genomes release 40 |
| *Bacillus murimartini* | GCA_001274705.1 | *Bacillus murimartini* | Ensembl Genomes release 40 |
| *Bacillus paralicheniformis* | GCA_000746885.1 | *Bacillus paralicheniformis* | Ensembl Genomes release 40 |
| *Bacillus pumilus* | GCA_000590455.1 | *Bacillus pumilus* | Ensembl Genomes release 40 |
| *Bacillus subtilis* subsp. *inaquosorum* | GCA_000332645.1 | *Bacillus subtilis* subsp. *inaquosorum* | Ensembl Genomes release 40 |
| *Bacillus subtilis* subsp. *spizizenii* | GCA_000816805.1 | *Bacillus subtilis* subsp. *spizizenii* | Ensembl Genomes release 40 |
| *Bacillus tequilensis* | GCA_001278955.1 | *Bacillus tequilensis* | Ensembl Genomes release 40 |
| *Bacillus thuringiensis* | GCA_000600315.1 | *Bacillus thuringiensis* | Ensembl Genomes release 40 |
| *Clostridium perfringens* | CP019468.1 | *Clostridium perfringens* | ENA Release 137 |
| *Geobacillus icigianus* | GCA_001587475.1 | *Geobacillus* sp b4113 201601 | Ensembl Genomes release 40 |
| *Geobacillus thermodenitrificans* | GCA_001587475.1 | *Geobacillus thermodenitrificans* | Ensembl Genomes release 40 |
| *Jeotgalibacillus marinus* | GCA_001274925.1 | *Jeotgalibacillus marinus* | Ensembl Genomes release 40 |
| *Listeria monocytogenes* | GCA_000382925.1 | *Listeria monocytogenes* | Ensembl Genomes release 40 |
| *Staphylococcus aureus* | GCA_000597965.1 | *Staphylococcus aureus* | Ensembl Genomes release 40 |
| *Staphylococcus delphini* | PRJEA8701 | *Staphylococcus delphini* | ENA Release 137 |
| *Staphylococcus epidermidis* | GCA_000759555.1 | *Staphylococcus epidermidis* | Ensembl Genomes release 40 |
| *Staphylococcus equorum* | GCA_001747895.1 | *Staphylococcus equorum* | Ensembl Genomes release 40 |
| *Staphylococcus haemolyticus* | GCA_000009865.1 | *Staphylococcus haemolyticus* | Ensembl Genomes release 40 |
| *Staphylococcus lugdunensis* | GCA_000247225.2 | *Staphylococcus lugdunensis* | Ensembl Genomes release 40 |
| *Staphylococcus pseudintermedius* | GCA_001622965.1 | *Staphylococcus pseudintermedius* | Ensembl Genomes release 40 |

**Table S4:** Non-dimensional system parameters and their values

| Descriptor | Symbol | Non-Dimensional Value |
| --- | --- | --- |
| Biomass density (assumed fixed) | $B$ | 1 |
| Initial iron concentration in the agar | $F_0$ | 1* |
| Iron level resulting in half maximal growth rate | $F_h$ | 1 |
| Biomass material constant | $\lambda$ | 1 |
| Growth response to change in iron level | $m$ | 3.5 |
| Rate constant for pulcherriminic acid production wild type | $k_b$ | 0.17 |
| Rate constant for pulcherriminic acid minus strains | $k_b$ | 0 |
| Rate constant for iron chelation (pulcherrimin production) | $k_p$ | $0.6 k_b$ |
| Rate constant for iron utilisation due to biomass production | $k_f$ | 0.17 |
| Pulcherriminic acid diffusion rate constant | $D_A$ | 10 |
| Iron diffusion rate constant | $D_F$ | $10^{-4} D_A$ |
| Pulcherrimin diffusion rate constant | $D_P$ | $10^{-4} D_A$ |

\*The value of initial free iron was varied as illustrated in Figure S6.

**Figure S1:**
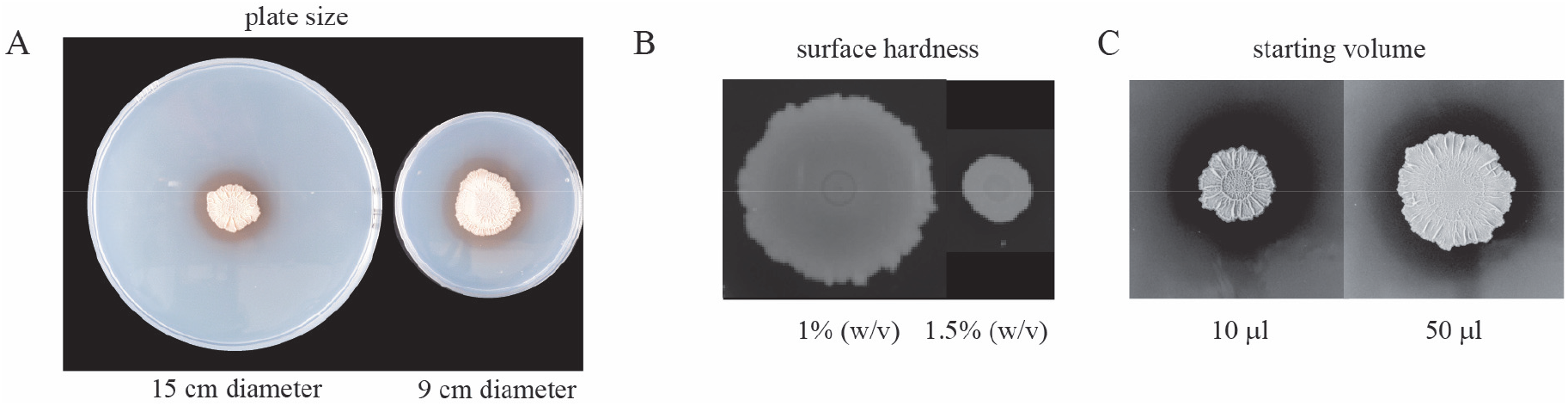
Environmental conditions influence the area colonised by the biofilm. **(A)** NCIB 3610 biofilms formed on 15cm and 9 cm diameter plates; **(B)** NCIB 3610 biofilms formed on 9 cm diameter plates solidified with either 1% or 1.5% w/v agar; **(C)** NCIB 3610 biofilms formed on 9 cm diameter plates using different inoculation volume that contained the same number of cells.

**Figure S2:**
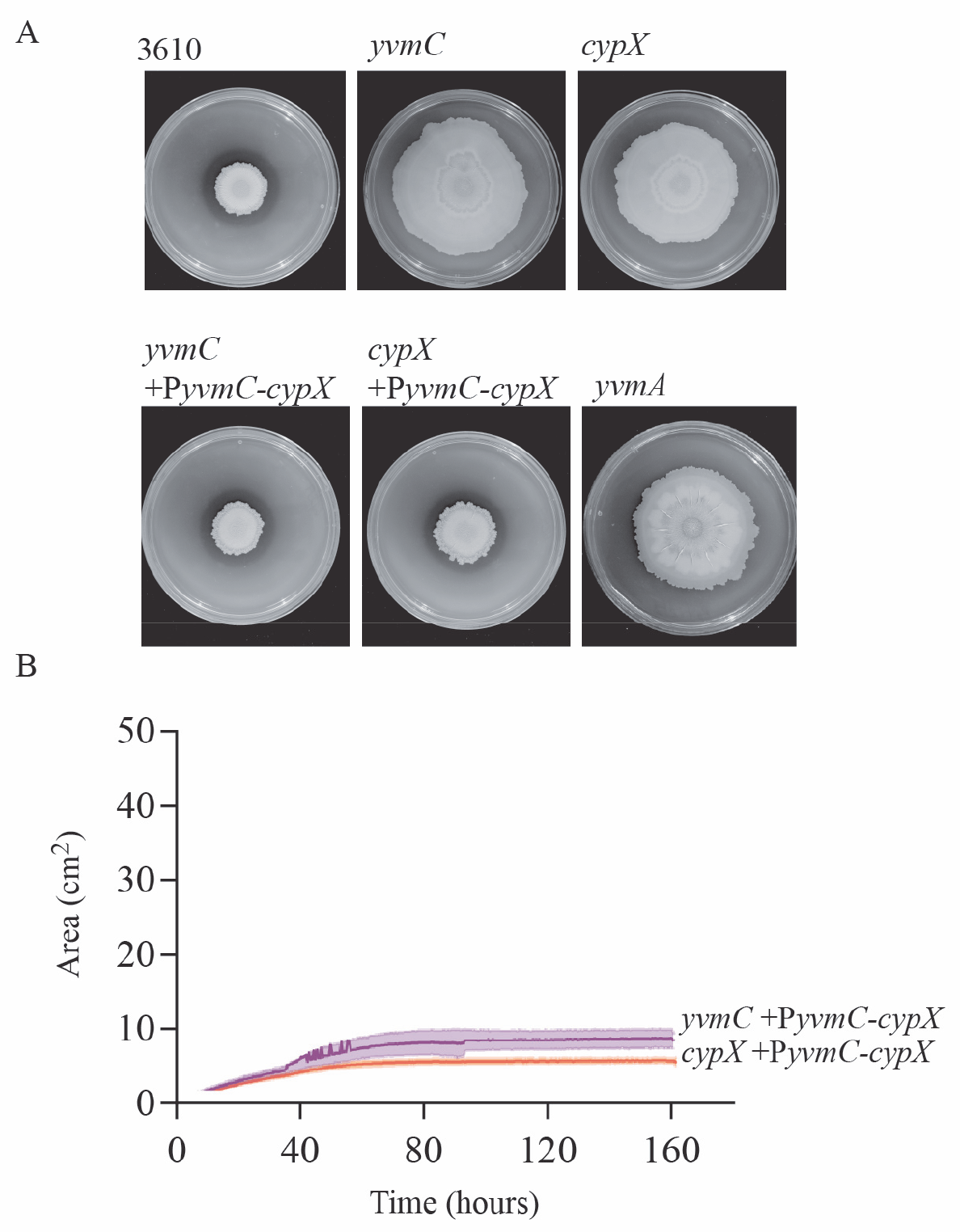
Genetic complementation of the pulcherriminic acid deficient strains. **(A)** Representative images of biofilms formed by strains NCIB 3610, *cypX* (NRS5532), *yvmC* (NRS5533), *yvmC amyE:PyvmC-yvmC-cypX* (NRS6240), *cypX amyE:PyvmC-yvmC-cypX* (NRS6239) and *yvmA* (NRS6248) after 120 hours growth at 30°C on MSgg agar containing 50 μM FeCl_3_.; **(B)** The area occupied by the biofilm formed by strains *yvmC amyE:PyvmC-yvmC-cypX* (NRS6240) and *cypX amyE:PyvmC-yvmC-cypX* (NRS6239) on the 9 cm diameter petri dish was calculated and plotted. The solid lines represent the average of 3 biological repeats and the shaded areas the standard error of the mean.

**Figure S3:**
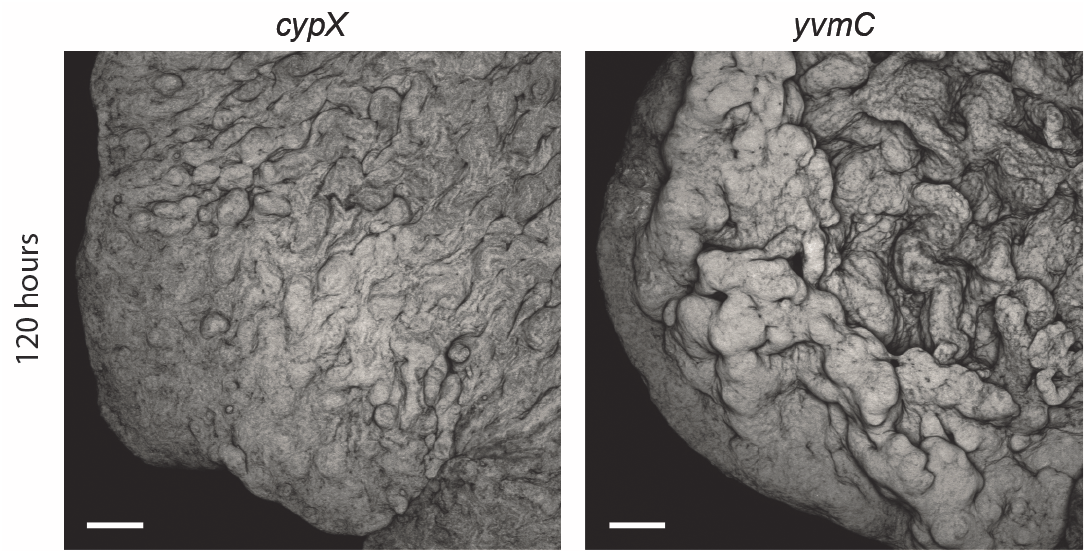
Confocal analysis of the *cypX* and *yvmC* mutants at 120 hours. Confocal microscopy of the biofilm edge at 120 hours for the *cypX* (NRS5532) and *yvmC* (NRS5533) deletion strains. In each case the strains were mixed with an isogenic variant carrying a constitutively expressed copy of *gfp*. This allowed detection of a fraction of the cells in the biomass by confocal microscopy. The images shown are projections of the acquired z-stacks and the scale bars represent 100 μm.

**Figure S4:**
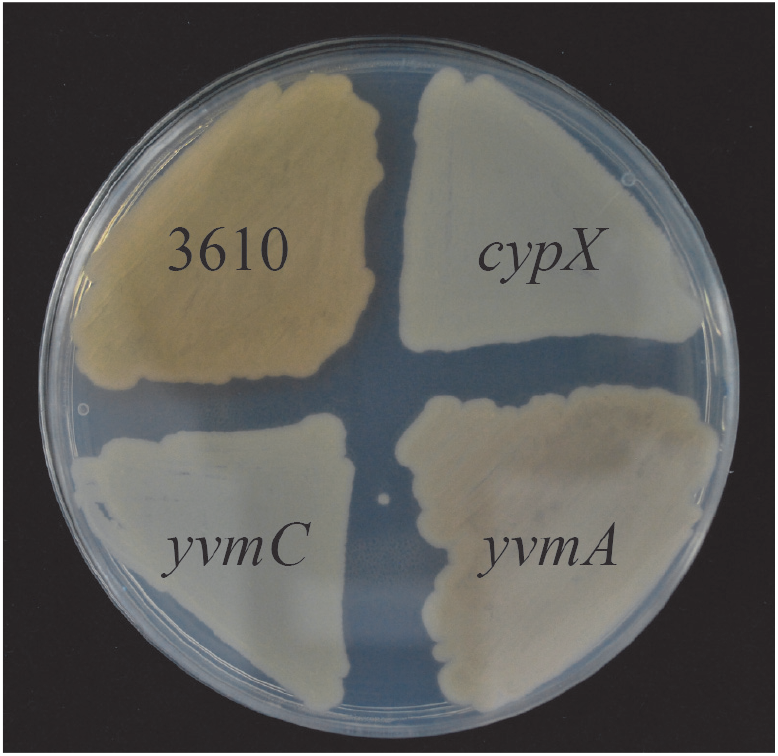
YvmA transports pulcherriminic acid to the extracellular environment. NCIB 3610, *yvmC* (NRS5533), *cypX*(NRS5532) and *yvmA* (NRS6248) strains grown on an MSgg agar plate for 24 hours at 37°C prior to imaging.

**Figure S5:**
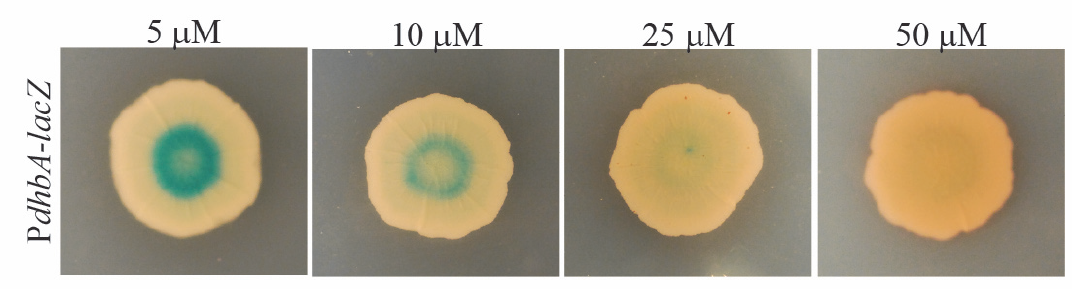
*PdhbA* transcription monitored *in situ* during biofilm formation. Images of 3610 containing the *PdbhA-lacZ* transcriptional reporter fusion after growth at 30°C on MSgg agar in 9 cm diameter petri dishes containing 120 μg ml^−1^ 5-bromo-4-chloro-3-indolyl-β-D-galactopyranoside (X-gal) for 48 hours. The starting level of FeCl_3_ in the growth medium was 5, 10, 25 and 50 μM as indicated. The image shown is representative of three biological repeats.

**Figure S6:**
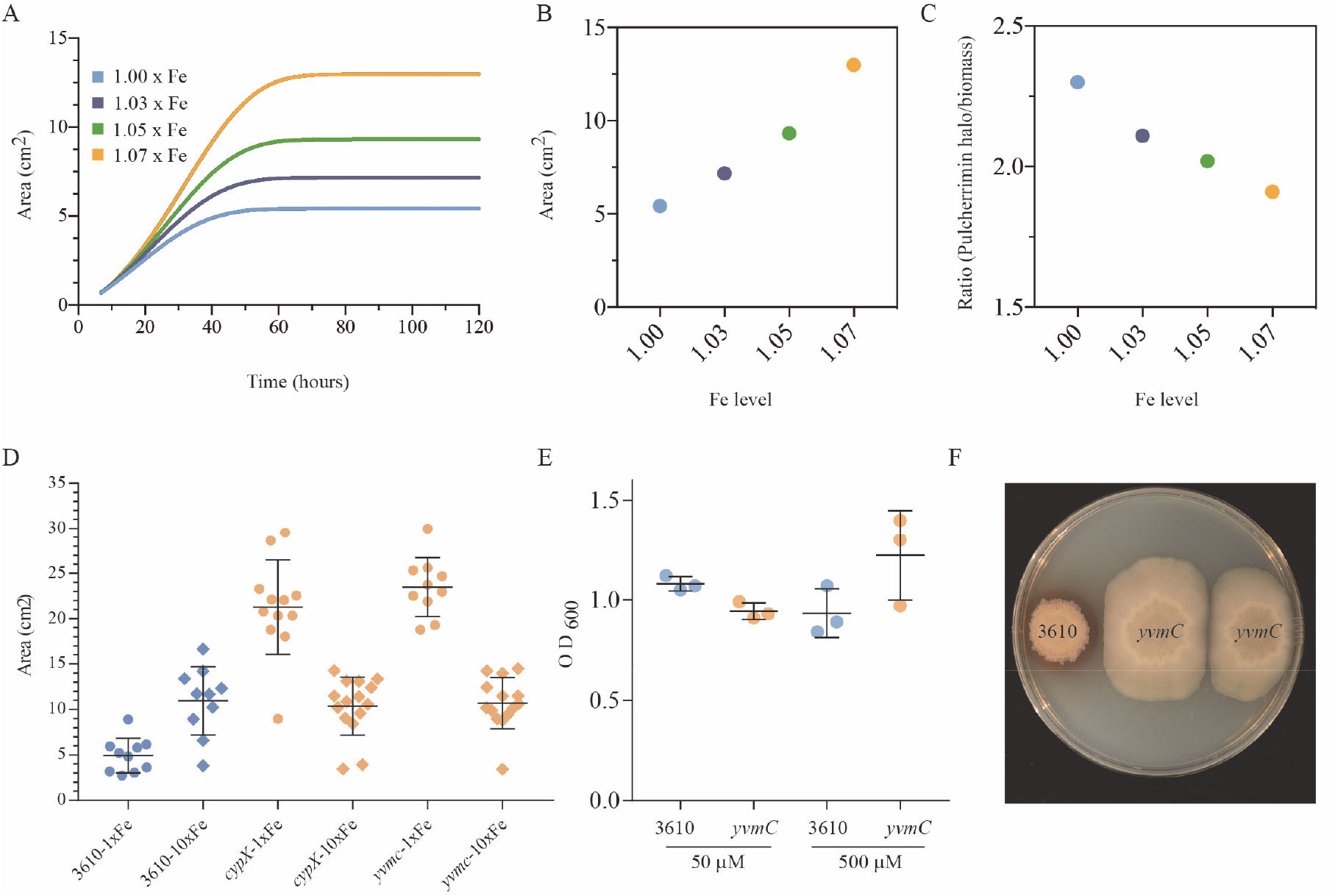
Increasing the level of iron in the external environment transiently overcomes self-restriction of growth. **(A)** Model prediction for the biofilm surface area over time for different values of the initial iron concentration. **(B)** Terminal biofilm surface area for (A) versus the initial iron concentration. **(C)** Ratio of the Pulcherrimin halo radius to that of the biofilm footprint as given in **(B)**, versus the initial iron concentration. The boundary of the biomass is shown as a dotted line. **(D)** NCIB 3610 and the *cypX* (NRS5532) and *yvmC* (NRS5533) deletion strains were grown at 30°C on MSgg agar containing either 50 μM FeCl_3_ (1x) or 500 μM FeCl_3_ (10x) for 120 hours prior to calculation of the area occupied. The bar represents the average of the biological repeats that are presented as individual points, the error bars are the standard error of the mean; **(E)** NCIB 3610 and the *cypX* (NRS5532) and *yvmC* (NRS5533) deletion strains were grown for 10 hours in liquid MSgg medium containing either 50 μM FeCl_3_ (1x) or 500 μM FeCl_3_ (10x) prior to measurement of the cell density using OD_600_ measurements. The bar represents the average of the biological repeats that are presented as individual points, the error bars are the standard error of the mean; **(F)** Biofilms of NCIB 3610 and *yvmC* (NRS5533) were inoculated onto a 15 cm diameter MSgg agar (1.5% w/v agar) plate and incubated at 30°C for 120 hours prior to imaging.

